# Inhibition of PAK2 in endothelial cells suppresses tumor angiogenesis and promotes immune sensitization through CXCL10

**DOI:** 10.1101/2024.04.05.588280

**Authors:** Jeanne Corriveau, Pascale Monot, Chantal Delisle, Marie-Anne Goyette, Camille Gasse, Yassine El Bakkouri, Trang Hoang, Jean-François Côté, Jean-Philippe Gratton

## Abstract

Tumor angiogenesis is induced by the secretion of pro-angiogenic factors and results in a disorganized tumor vasculature. The p21-activated kinase 2 (PAK2) is involved in endothelial cell (EC) migration, an essential step of angiogenesis. However, the involvement of PAK2 in tumor angiogenesis remains ill-defined. Here, we show that, during orthotopic tumor growth, the specific deletion of PAK2 in ECs reduces tumor size and angiogenesis. In addition, endothelial-specific loss of PAK2 was found to normalize the remaining tumor blood vessels, favoring innate immune cells infiltration. Importantly, we uncovered a role for PAK2 in regulating chemokine expression, notably CXCL10. Secretion of CXCL10 from ECs is enhanced following PAK2 depletion and its expression is essential for the inhibitory effects of PAK2 silencing on EC spouting. Furthermore, neutralization of CXCL10 in mice reversed the effects induced by the deletion of PAK2 on tumor vasculature and immune composition of tumors. Together, our findings identify endothelial PAK2 as a potential target to reduce tumor angiogenesis and reprogram ECs to promote immune cell infiltration within tumors through the expression of CXCL10.

## INTRODUCTION

The tumor microenvironment (TME), the ecosystem in which tumors develop, plays an active role in the pathogenesis of cancer as it influences both neoplasia progression and the response to therapy ^1^. In addition to the primary cancer cells that form the bulk of the tumor, a wide variety of non-cancerous cells, including endothelial cells (ECs), fibroblasts, and immune cells compose the TME. ECs contribute to tumor angiogenesis, the formation of new blood vessels that supply nutrients and oxygen to the tumor ^2^. Tumor vascularization is essential for the growth of solid tumors as non-vascularized tumors do not grow more than a few millimeters in diameters ^2,3^. Indeed, as a tumor grows, cancer cells become hypoxic, leading to their reprogramming to express pro-angiogenic factors, such as hypoxia-inducible factors (HIFs), vascular endothelial growth factor (VEGF), platelet-derived growth factor (PDGF), angiopoietins and other cytokines, that collectively operate to initiate neo-angiogenesis to meet their metabolic needs ^4,5^. However, due to the unbalanced increase of pro-angiogenic factors, tumor blood vessels exhibit an immature phenotype compared to vessels in normal tissues ^6,7^. Tumor vessels are excessively branched, chaotic and leaky, as they lack pericyte and smooth muscle coverage, and possess an inconsistent basement membrane with irregular endothelial lining. This phenotype leads to inconsistent perfusion and a hypoxic tumor state, contributing to the sustained secretion of pro-angiogenic factors ^6^.

Hypoxia also contributes to the chronic inflammatory state of tumors, which is an essential component of the TME and one of the hallmarks of cancer ^8^. Hypoxia leads to the upregulation of pro-inflammatory cytokines and C-C and C-X-C chemokines that promote the recruitment of immune cells to the hypoxic regions of the tumor ^9^. These recruited cells directly contribute to the accentuation of tumor blood vessels abnormalities by releasing pro-angiogenic factors in the TME ^10^. Also, inflammatory cytokines and chemokines can promote the recruitment of immunosuppressive cells, such as tumor-associated macrophages and regulatory T cells (Tregs), and inhibit the function of cytotoxic T cells and natural killer (NK) cells, that collectively contributes to immune suppression and immune evasion ^11^. In addition their effects on chemotaxis, C-C and C-X-C chemokines play important roles in tumor angiogenesis, having either angiogenic or angiostatic effects ^12^. For instance, CXCL8, CXCL12, CCL2, and CCL5 promote the migration and proliferation of ECs, while CXCL4, CXCL9, CXCL10 and CXCL14 inhibit these processes ^12,13^. Although its expression has been associated with tumor progression in certain types of cancer, CXCL10 expression in the TME is generally associated with anti-tumor effects through immunogenic and angiostatic activities ^14^. Combining anti-angiogenic agents with immunotherapy has been proposed to improve treatment efficacy ^15^. However, anti-angiogenic agents, mostly targeting the VEGF-signaling pathways, have encountered therapeutic resistance in the clinic, partly due to the activation of alternative pro-angiogenic pathways ^16^.

p21-activated kinases (PAKs) are serine/threonine kinase best known to be effectors of the small Rho GTPases of the Rac- and Cdc42-subfamillies. The PAK family is divided into Group I (PAK1, 2, 3) and Group II (PAK4, 5, 6), and is involved in various cellular processes, including cytoskeletal remodeling, cell motility, proliferation and survival ^17^. Among Group I PAKs, PAK2 is ubiquitously expressed, whereas the expressions of PAK1 and PAK3 exhibit tissue specificity ^18^. PAK1, 2 and 4 are expressed in vascular tissues, with PAK2 being the dominant PAK isoform expressed in ECs ^18,19^. PAK1-deficient mice are viable and fertile with minor immune, neuronal and glucose homeostasis defects ^20^. PAK4-deficient mice show heart and neuronal defects and lack vascular branching leading to embryonic lethality, but specific deletion of *PAK4* in ECs of adult mice did not affect normal endothelial functions ^20,21^. In contrast, PAK2 knockout (KO) mice, as well as mice with endothelial-specific deletion of PAK2, are embryonic lethal due to vascular defects including impaired vascular remodeling and altered barrier function ^20,22,23^. In adult mice, induction of global PAK2 genetic deletion is also lethal and the induction of endothelial-specific depletion results in increased vascular permeability in various tissues ^23^. In ECs, PAK2 activation is necessary for VEGF- and angiopoietin-1-stimulated migration and angiogenic sprouting ^23–25^. Interestingly, endothelial-specific deletion of PAK4 in mice inhibited angiogenesis and growth of glioblastoma tumors ^21^. Surprisingly, despite being the dominant isoform expressed in ECs, the role of PAK2 in tumor angiogenesis remains ill-defined.

Here, we report that induction of endothelial-specific deletion of PAK2 during the growth of orthotopic tumors in mice results in reduced tumor size and angiogenesis and induces the normalization of tumor blood vessels. Notably, deletion of PAK2 in tumor ECs increases the presence of innate immune cells into tumors, in particular NK and dendritic cells (DCs), and results in augmented expression of chemokines, including CXCL10. Furthermore, secretion of CXCL10 from ECs is augmented following PAK2 silencing. Importantly, neutralization of CXCL10 with blocking antibodies reverses the anti-tumor effects induced by PAK2 deletion. Together, our findings show that the endothelial-specific deletion of PAK2 in tumors inhibits tumor angiogenesis and reprograms ECs to promote, in a CXCL10-dependent manner, immune cell infiltration limiting tumor progression.

## RESULTS

### Conditional deletion of PAK2 in tumor ECs suppresses tumor angiogenesis and growth

The goal of this study is to define the contributions of PAK2 to tumor angiogenesis. Since endothelial-specific loss of PAK2 induces lethality during mouse embryonic development ^23^, we generated a mouse line that allowed us to induce endothelial *Pak2* gene deletion in adult mice upon tamoxifen treatment to determine the functions of PAK2 in tumor blood vessels. Pak2^fl/fl^ mice ^23^, were crossed with *Pdgfb*-iCreER mice ^26^ to generate *Pdgfb*-iCreER;Pak2^fl/fl^ mice which were viable and fertile with no obvious phenotypes. The *Pdgfb*-iCreER transgenic mouse model is particularly suited to study tumor angiogenesis since the *Pdgfb* promotor drives the expression of the CreER preferentially in ECs of growing and immature blood vessels ^26,27^. *Pdgfb*-iCreER;Pak2^fl/fl^ mice were genotyped by PCR (Fig. S1A). In this model, tamoxifen treatment of *Pdgfb*-iCreER;Pak2^fl/fl^ mice will result in specific deletion of *Pak2* in ECs of mice (KO-PAK2^EC^). Pak2^fl/fl^ (CT-PAK2^EC^) or *Pdgfb*-iCreER mice (CT-CreER) treated with tamoxifen or *Pdgfb*-iCreER;Pak2^fl/fl^ treated with a placebo (PCB-PAK2^EC^) were used as controls (Fig. 1A). To verify the efficiency of PAK2 protein loss *ex vivo* in ECs, mouse lung ECs (MLECs) were isolated from *Pdgfb*-iCreER;Pak2^fl/fl^ mice and treated with increasing concentrations of 4-hydroxytamoxifen (OH-tamoxifen) for 24 h *ex vivo*. As predicted, MLECs isolated from *Pdgfb*-iCreER;Pak2^fl/fl^ mice and treated with OH-tamoxifen showed a significant reduction in PAK2 protein levels (Fig. S1B).

**Figure 1.**
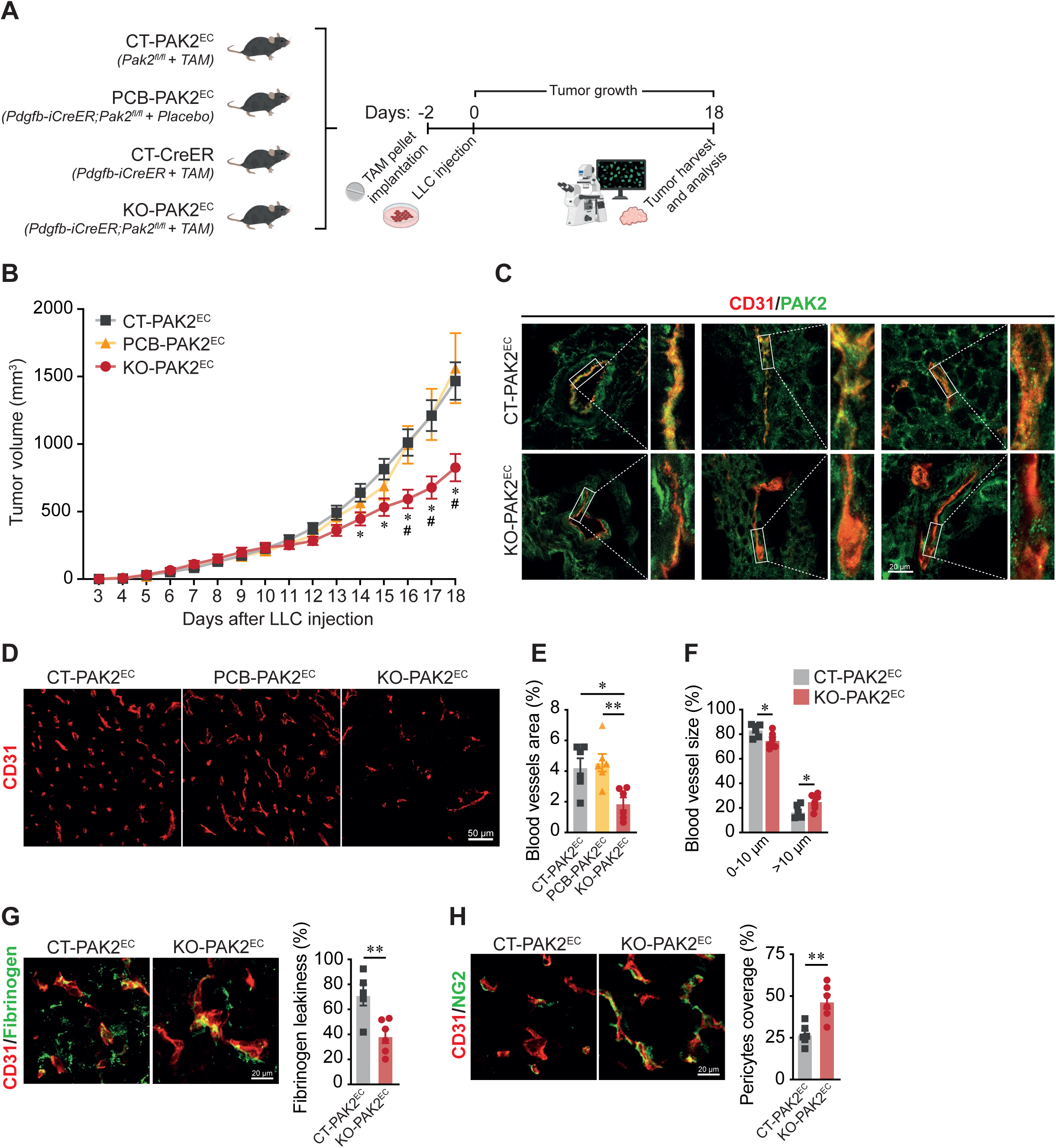
Deletion of PAK2 in tumor ECs inhibits tumor angiogenesis and growth, and normalizes the tumor vasculature. **(A)** Schematic illustration of cohort experiments performed on 6 to 8 weeks old mice. Mice were treated with placebo or tamoxifen (TAM; slow-release pellet, 5 mg/21 days s.c.) and injected with 1.5x10^6^ Lewis lung carcinoma (LLC) cells (s.c.). Created with Biorender.com. **(B)** Growth curves of LLC tumors implanted on CT-PAK2^EC^ (n=31 mice), PCB-PAK2^EC^ (n=14 mice) and KO-PAK2^EC^ mice (n=31 mice). *p<0.05 between CT-PAK2^EC^ and KO-PAK2^EC^, #p<0.05 between PCB-PAK2^EC^ and KO-PAK2^EC^. **(C)** Visualization of the endothelial-cell specific deletion of PAK2 in tumor sections. Scale = 20 μm. **(D)** Representative images of CD31^+^ staining in tumors sections. Scale = 50 μm. **(E)** Quantification of CD31^+^ vascular area in tumors (CD31^+^ area/total area) (n=6 tumors). **(F)** Quantification of diameters of CD31^+^ vascular structures in tumors (n=6 tumors). **(G)** Quantification of NG2^+^ pericyte coverage of blood vessels in tumors (NG2^+^ area /CD31^+^ area) (n=6 tumors). Scale = 20 μm. **(H)** Quantification of Fibrinogen leakage from blood vessels in tumors (Fibrinogen^+^ area/CD31^+^ area) (n=5-6 tumors). Scale = 20 μm. All data are represented as mean ± SEM. *p<0.05, **p<0.01.

In order to determine if the loss of PAK2 in ECs affects tumor angiogenesis, Lewis Lung Carcinoma (LLC) tumor cells were injected subcutaneously into the flank of KO-PAK2^EC^, CT-PAK2^EC^, CT-CreER or PCB-PAK2^EC^ mice. Tumors were allowed to grow for a maximum of 18 days and tumor size was measured daily (Fig. 1A). The growth of tumors on KO-PAK2^EC^ mice was significantly reduced compared to tumors on CT-PAK2^EC^, CT-CreER or PCB-PAK2^EC^ mice (Fig. 1B; Fig. S1C). The endothelial-specific loss of PAK2 within the tumor vasculature of KO-PAK2^EC^ mice was confirmed by CD31 and PAK2 immunofluorescence staining of tumor tissue sections. Tumor isolated from KO-PAK2^EC^ showed absence of CD31 and PAK2 colocalization in contrast to staining in CT-PAK2^EC^ tumors, while PAK2 staining outside tumor ECs remained unaffected (Fig. 1C). To assess the effect of endothelial PAK2 loss on tumoral vascularization, the CD31-positive area was quantified by immunofluorescence staining of tumor tissue sections. KO-PAK2^EC^ tumors had less CD31 staining, an EC marker, compared to tumors implanted on PCB-PAK2^EC^ or on CT-PAK2^EC^ (Fig. 1D, 1E). Due to the lack of significant differences in tumor growth or levels of intra-tumoral vascularization between all control groups, we elected to use CT-PAK2^EC^ mice (Pak2^fl/fl^ + Tamoxifen) as our main control group for further experiments. Overall, these results show that the endothelial-specific deletion of PAK2 reduces both tumor growth and tumor vascularization, demonstrating an essential role for endothelial PAK2 in the TME in determining tumor progression and angiogenesis.

An immature and disorganized vasculature is a hallmark of tumor blood vessels ^6^. We, therefore, investigated if the loss of PAK2 expression in tumor ECs affects the structural and functional states of tumor blood vessels. Despite the overall reduced tumoral vascularization, the blood vessels found in KO-PAK2^EC^ tumors were of larger caliber, showing more vascular structures of wider diameter (> 10 μm), than the blood vessels found in CT-PAK2^EC^ tumors (Fig. 1D, 1F). KO-PAK2^EC^ tumor blood vessels also possess higher pericyte coverage compared to control tumors, with increased levels of the pericyte marker NG2 relative to CD31 levels in tumors (Fig. 1G). Since these results suggested that the blood vessels in KO-PAK2^EC^ tumors were better supported by pericytes and more stable, we examined the extent of their leakiness to fibrinogen. KO-PAK2^EC^ tumors had a decreased extravascular deposition of fibrinogen compared to CT-PAK2^EC^ tumors (Fig. 1H), indicating that KO-PAK2^EC^ tumor blood vessels are less permeable to macromolecule extravasation. Altogether, these results show that the deletion of PAK2 in ECs suppresses tumor angiogenesis, reducing both tumor growth and vascularization, and leads to normalization of tumor blood vessels.

### KO-PAK2^EC^ tumors express an immune response gene signature

To gain insights into the consequences of endothelial PAK2 deletion on tumors, we performed RNA-sequencing (RNA-seq) analysis of three KO-PAK2^EC^ and three CT-PAK2^EC^ tumors. Differential gene expression analyses identified 478 upregulated and 166 downregulated transcripts in KO-PAK2^EC^ tumors compared to CT-PAK2^EC^ tumors (Fig. 2A and Table S1). Differential gene expression of selected transcripts was confirmed by RT-qPCR (Fig. S2A). Gene ontology (GO) analysis centered on biological processes of the 644 differentially expressed genes (DEGs) in KO-PAK2^EC^ tumors identified several significant GO annotations (Table S2). Interestingly, many of the top 20 most significant GO annotations of DEGs in KO-PAK2^EC^ tumors were associated to an activation of the immune response, such as defense response (GO:0006952), innate immune response (GO:0045087), response to interferon-beta (GO:0035456), immune response (GO:0006955) and response to interferon-gamma (GO:0034341) (Fig. 2B). Gene set enrichment analysis (GSEA) confirmed that the immune response gene set (GO:0006955) was enriched in KO-PAK2^EC^ tumors compared to CT-PAK2^EC^ tumors (Fig. 2C). In contrast, the sprouting angiogenesis gene set (GO:0002040) was enriched in CT-PAK2^EC^ tumors, consistent with the reduction in vascularization levels we observed in KO-PAK2^EC^ tumors (Fig. 2D). Interestingly, *Esm1*, an endothelial tip cell and neoangiogenesis marker, is among the genes that contributed to the enrichment of this gene set and was downregulated in KO-PAK2^EC^ tumors ^28^. This suggest that the reduction of blood vessels observed in KO-PAK2^EC^ tumors is accompanied by a reduction in endothelial tip cells (Fig. 2D). Response to oxygen levels (GO:0070482) and response to hypoxia (GO:0001666) did not show significant enrichment (FDR>0.25) between groups (Fig. S2E-F). However, the expression of Hypoxia-inducible factor 1-alpha (*Hif1a*) was slightly upregulated, though not significant, in KO-PAK2^EC^ compared to CT-PAK2^EC^ tumors (Fold increase = 1.2, adjusted p-value = 0.06) (Table S1, Fig. S2F). This indicated that tumor hypoxia is not the main driving factor for the effects induced by endothelial PAK2 deletion. Regulation of programmed cell death (GO:0043067), inflammatory response (GO:0006954) and response to cytokine (GO:0034097) gene sets were enriched in KO-PAK2^EC^ tumors compared to CT-PAK2^EC^ tumors (Fig. S2B-D). These analyses strongly suggest that KO-PAK2^EC^ tumors are more immunologically active than CT-PAK2^EC^ tumors. Furthermore, analysis of all the significant GO annotations identified (Table S2) revealed that many of them were associated with the response to cytokine, adhesion and chemotaxis of immune cells (Fig. 2E). Leukocyte cell-cell adhesion (GO:0007159) and leukocyte chemotaxis (GO:0030595) gene sets were significantly enriched in KO-PAK2^EC^ tumors (Fig. 2F, 2G). Taken together, these analyses reveal that the conditional loss of PAK2 in ECs rewires the transcriptomic profile of KO-PAK2^EC^ tumors, resulting in a microenvironment that may promote the infiltration of immune cells.

**Figure 2.**
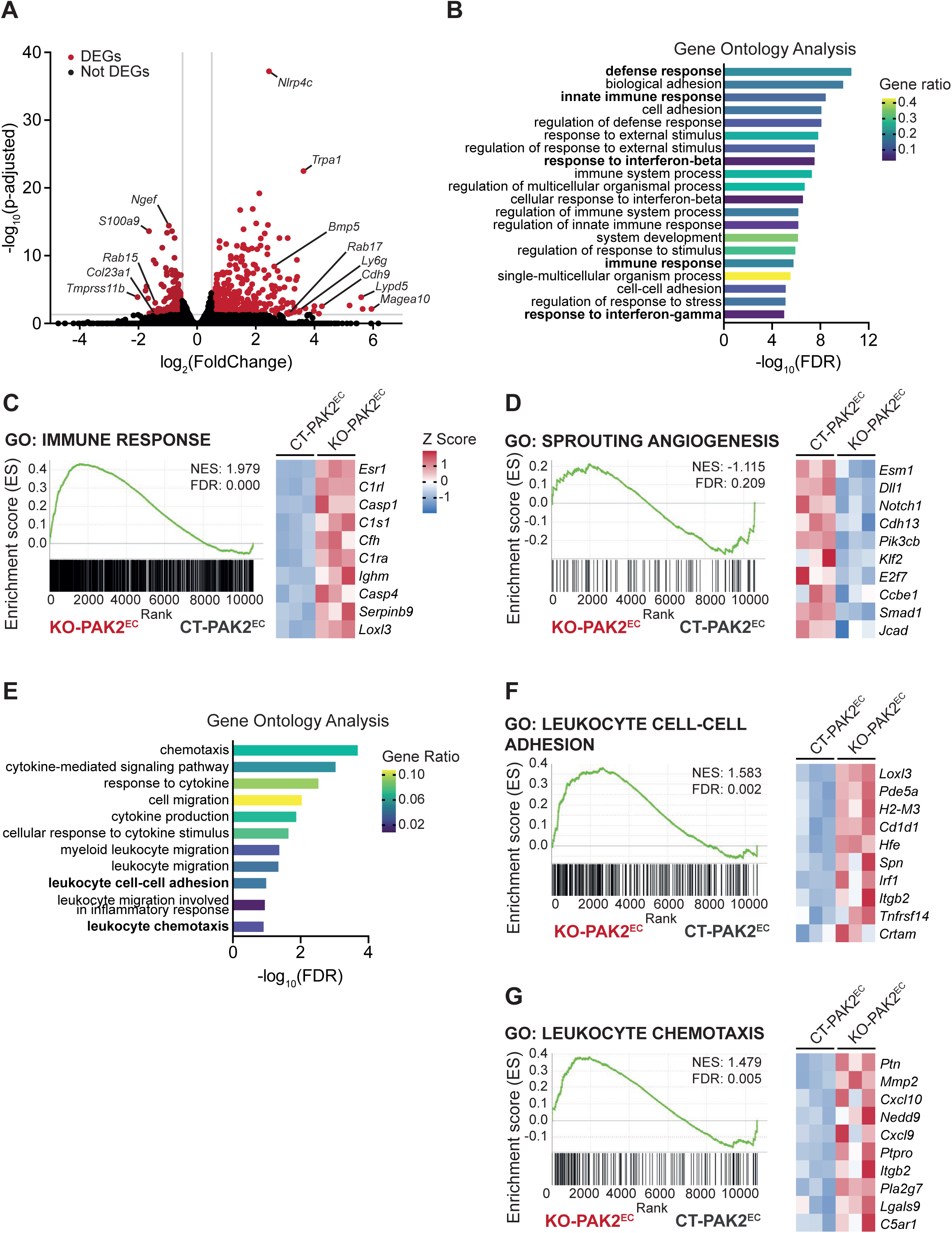
Endothelial-specific deletion of PAK2 in tumors transcriptionally reprograms the the tumor microenvironment. **(A)** Volcano plot showing DEGs in red circles (644 genes) between KO-PAK2^EC^ (n=3 tumors) and CT-PAK2^EC^ (n=3 tumors) from bulk RNA-seq. A threshold of a log_2_(FoldChange) >0.5 and <-0.5, and a p-adjusted value <0.05 was used to define DEGs. **(B)** Bar chart of the top 20 significant GO terms of DEGs submitted to the DAVID database and categorized using the biological process annotations. The x-axis indicates the statistical significance of the enrichment (-log_10_(FDR)). Color-coding indicates the gene ratio, referencing to the ratio of DEGs annotated to all genes annotated in the given GO term. **(C-D)** GSEA of GO terms (C) immune response (GO:0006955) and (D) sprouting angiogenesis (GO:0002040) of KO-PAK2^EC^ compared to CT-PAK2^EC^ tumor gene expression. Heatmaps represent the top 10 genes contributing to the normalized enrichment score (NES). **(E)** Bar chart of selected GO terms of DEGs submitted to the DAVID database and categorized using the biological process annotations related to cytokine response, cell adhesion and cell chemotaxis. The x-axis indicates the statistical significance of the enrichment (-log_10_(FDR)). Color-coding indicates the gene ratio, referencing to the ratio of DEGs annotated in the given GO term to all genes annotated in the GO term. **(F-G)** GSEA of GO terms (F) leukocyte cell-cell adhesion (GO:0007159) and (G) leukocyte chemotaxis (GO:0030595) of KO-PAK2^EC^ compared to CT-PAK2^EC^ tumor gene expression. Heatmaps represent the top 10 genes contributing to the NES. The scale for Z scores in the heatmaps of panels D, F and G is shown in panel C.

### Deletion of PAK2 in tumoral ECs enhances innate immune cell infiltration

To assess the possibility that endothelial deletion of PAK2 triggers an immune response in tumors, we determined the immune cell profiles of KO-PAK2^EC^ and CT-PAK2^EC^ tumors by multiparametric flow cytometry analysis. These analyses showed a significant increase in the presence and/or infiltration of DCs, migratory CD103^+^ DCs, NK cells and activated CD44^+^ NK cells, all innate immune cell types, in KO-PAK2^EC^ tumors (Fig. 3A). We next investigated the localization of these immune cells within the tumor by immunohistochemical staining. Hematoxylin counterstain was used to determine core and periphery regions (Fig. S3A). Immunohistochemical staining of CD11c and NK1.1 showed that KO-PAK2^EC^ tumors had more DCs and NK cells in the core regions of tumors (Fig. 3B, 3D) while no significant difference was observed at the periphery of tumors (Fig. 3C, 3E). No significant difference in the content of adaptative immune cells was observed by flow cytometry analysis in KO-PAK2^EC^ and CT-PAK2^EC^ tumors (Fig. 3A). However, immunostaining of KO-PAK2^EC^ tumors revealed that they had less CD4^+^ T cells at their periphery than CT-PAK2^EC^ tumors, but no difference was observed in the core regions of tumors (Fig. S3B, 3C). No difference was observed for CD8^+^ T cells in the core or periphery of tumors (Fig. S3D, 3E).

**Figure 3.**
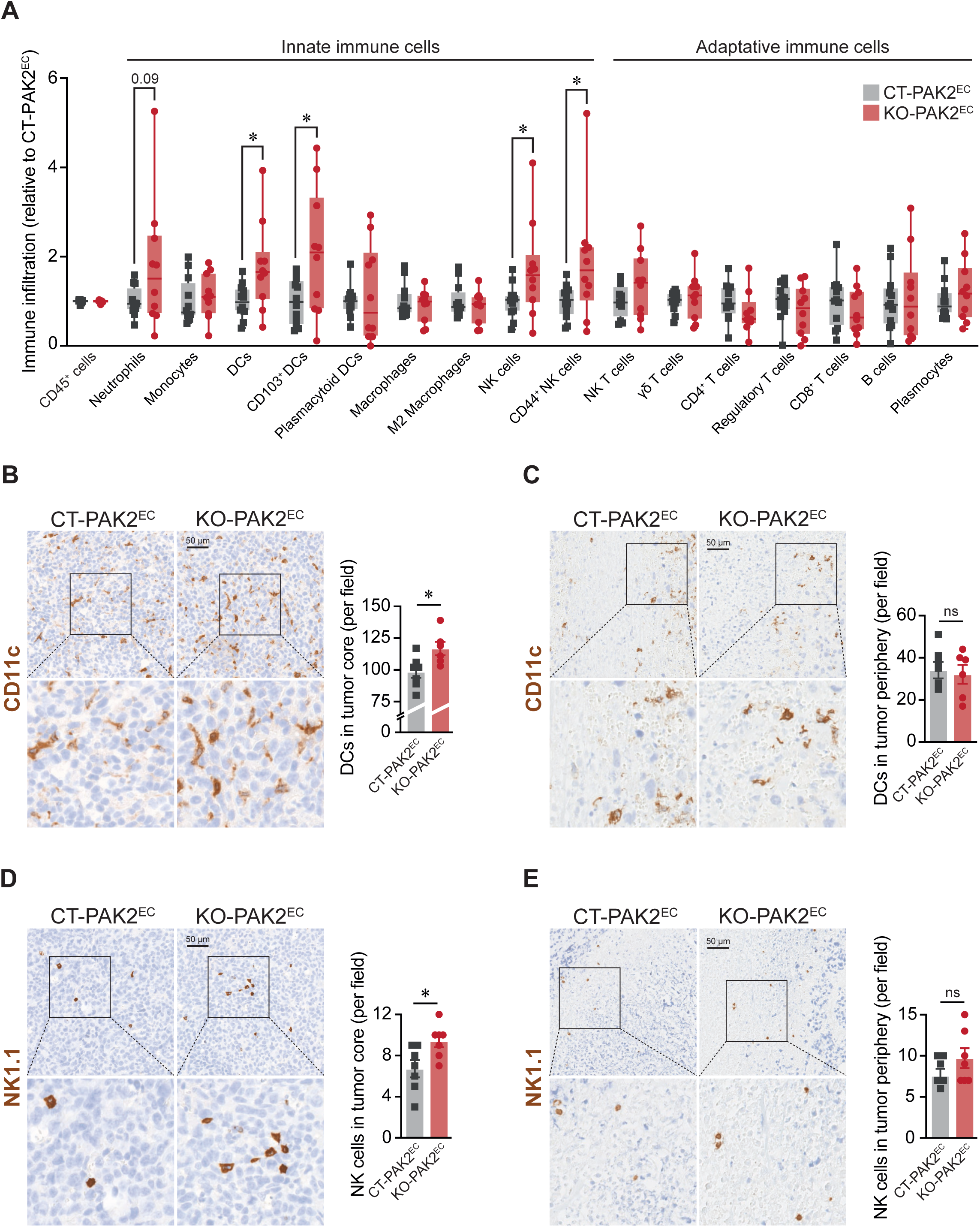
Deletion of PAK2 in ECs of tumors promotes innate immune cell infiltration. **(A)** Flow cytometry analysis of immune cell infiltration into KO-PAK2^EC^ tumors relative to CT-PAK2^EC^ tumors (n=10-12 tumors). DCs (CD45^+^I-A/I-E^+^CD11c^+^), migratory CD103^+^ DCs, NK cells (CD45^+^CD8^-^CD4^-^NK1.1^+^) and activated CD44+ NK cells are enriched in KO-PAK2^EC^. The number of immune cells for each population was normalized to the number of total live cells in each tumor. Combined data of three independent experiments are shown and represented as median ± minimal to maximal values. *p<0.05. **(B-E)** Immunohistochemistry analysis of KO-PAK2^EC^ and CT-PAK2^EC^ tumors. Quantification of DCs (CD11c^+^) in the (B) core regions and (C) periphery of tumors (n=5-6 tumors). Quantification of NK cells (NK1.1^+^) in the (D) core regions and (E) periphery of tumors (n=5-6 tumors). Data are represented as mean ± SEM. *p<0.05.

To determine if the immune transcriptomic changes observed in KO-PAK2^EC^ tumors are consistent with increased infiltration of innate immune cells, we further interrogated the DEGs between KO-PAK2^EC^ and CT-PAK2^EC^ tumors that were involved in the immune response gene set (GO:0006955) (Fig. S2G). Analysis of these 108 DEGs using Gene ontology centered on biological process revealed that the GO term innate immune response (GO:0045087) was more significantly enriched than the GO term adaptative immune response (GO:0002250) (Fig. S2H). In addition, pathway analysis using the KEGG database revealed that these genes were also enriched in pathways mostly involved in innate immune response such as NOD-like receptor signaling, C-type lectin receptor signaling and cytosolic DNA-sensing pathways, and pro-inflammatory cytokine responses, including IL-17 signaling, TNF signaling and NF-kappa B signaling pathways (Fig. S2I). Altogether, these results demonstrate that KO-PAK2^EC^ tumors had increased innate immune cell infiltration, principally DCs and NK cells.

### Inhibition of PAK2 increases the expression of inflammatory genes and of the chemokine CXCL10 in ECs

To determine how PAK2 expression in ECs could affect the immune cell content of tumors, we compared the transcriptomes of control and PAK2-depleted human umbilical vein ECs (HUVECs). Bulk RNA-seq was carried out on HUVECs transfected with either control (siCT) or siRNA targeting PAK2 (siPAK2). The PAK2 protein expression level was downregulated by more than 85% in siPAK2 transfected cells (Fig. S4A). We identified 1521 DEGs (986 upregulated and 535 downregulated genes) in siPAK2 compared to siCT transfected HUVECs (Fig. 4A and Table S3). As anticipated, *PAK2* was found among the most downregulated DEGs (Fold decrease = -4.8). The downregulation of PAK2 did not significantly affect the differential expression of other PAK genes (*PAK1*, *3*, *4*, *5*, *6*) (Fig. S4B, S4C). Interestingly, among the top 20 upregulated DEGs in siPAK2 transfected HUVECs, most of them are induced by interferon signaling (such as *MX2*, *MX1*, *IFI44L*, *IFIT1* and *RSAD2*) and four of the top 20 genes were chemokines (*CXCL10* [Fold increase = 30.7], *CX3CL1*, *CXCL11* and *CXCL12*) (Fig. S4D and Table S3). Furthermore, Gene ontology analysis revealed that many significant GO annotations of DEGs in siPAK2 were associated to an activation of the immune response and inflammation, such as response to cytokine (GO: 0034097), immune system process (GO: 0002376) and inflammatory response (GO: 0006954) (Fig. 4B). PAK2 downregulation in ECs also induced enrichment of the GO terms leukocyte adhesion to vascular endothelial cells (GO:0061756) and antigen processing and presentation of peptide antigen (GO:0048002), supporting a pro-inflammatory and pro-immune endothelial state following PAK2 silencing (Fig. S4E-F).

**Figure 4.**
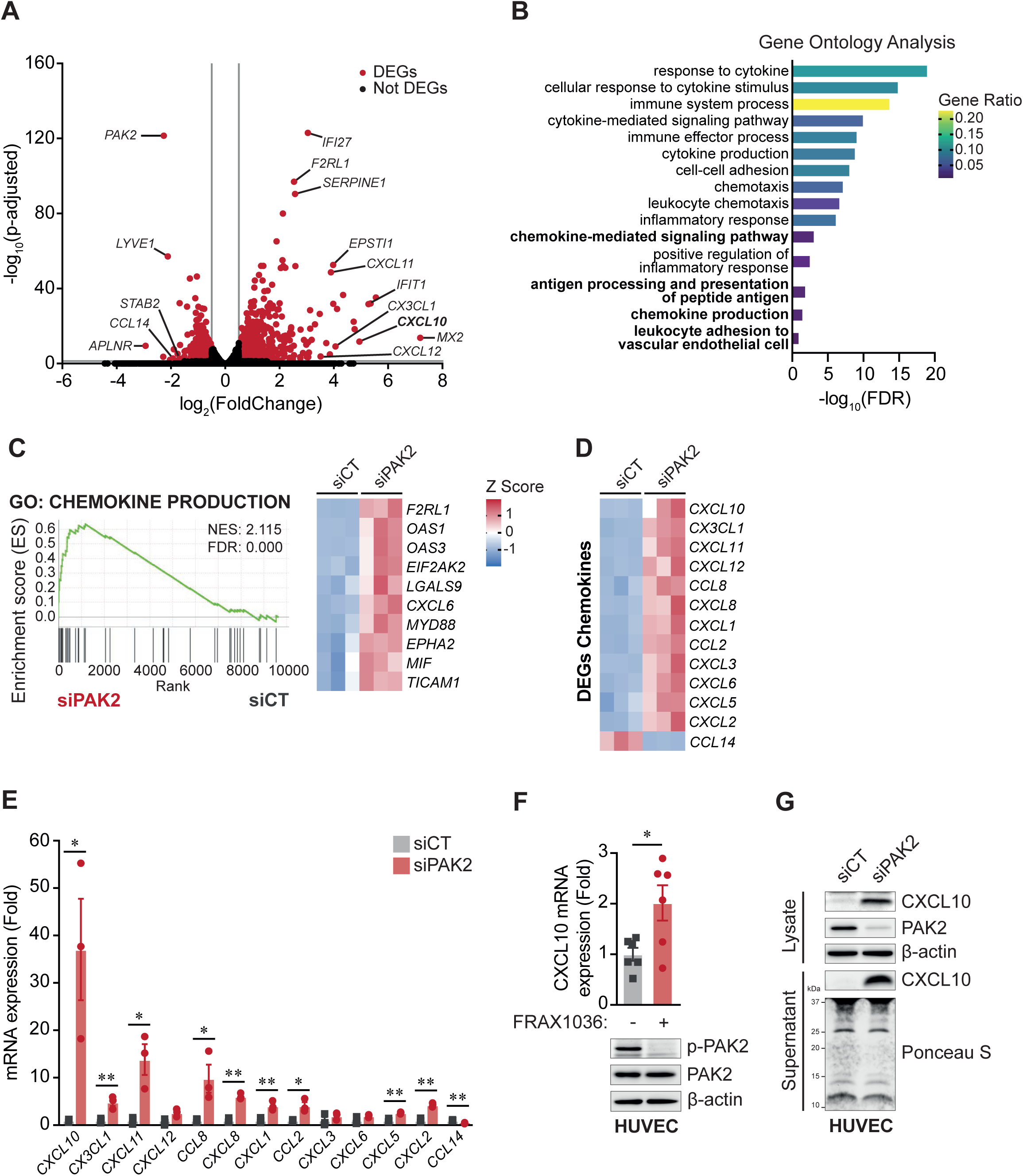
Downregulation of PAK2 in ECs increases the expression of inflammatory genes and CXCL10. **(A)** Volcano plot showing DEGs in red circles (1,521 genes) between siPAK2 (n=3) and siCT (n=3) transfected HUVECs from bulk RNA-seq. A threshold of a log_2_(FoldChange) >0.5 and <-0.5, and a p-adjusted value <0.05 was used to define DEGs. **(B)** Bar chart of selected GO terms of DEGs submitted to the DAVID database and categorized using the biological process annotations related to inflammatory and immune responses. The x-axis indicates the statistical significance of the enrichment (-log_10_(FDR)). Color-coding indicates the gene ratio, referencing to the ratio of DEGs annotated in the given GO term to all genes annotated in the GO term. **(C)** GSEA of GO term chemokine production (GO:0032602). Heatmaps represent the top 10 genes contributing to the normalized enrichment score (NES). **(D)** Heatmap of the chemokine DEGs in siPAK2 compared to siCT transfected HUVECs. The scale for Z scores in the heatmaps of panel D is shown in panel C. **(E)** Expression of chemokines genes measured by RT-qPCR, normalized to GAPDH housekeeping gene and expressed as fold change of siPAK2 relative to siCT transfected HUVECs. **(F)** Expression of CXCL10 gene measured by RT-qPCR, normalized to GAPDH housekeeping gene and expressed as fold change of FRAX1036 treated HUVECs (n=6) relative to non-treated HUVECs (n=6). PAK2 inhibition was monitored by Western blot analysis following FRAX1036 treatment (1 μM) for 7 h. β-actin served as loading control. **(G)** Protein expression of CXCL10 in cell lysate and cell supernatant of siPAK2 and siCT transfected HUVECs. PAK2 downregulation was monitored in siPAK2 transfected cells, and β-actin and Ponceau S staining served as loading control for cell lysate and cell supernatant respectively. All data are represented as mean ± SEM. *p<0.05. **p<0.01.

In addition, the presence of several chemokine-encoding genes among the top 20 genes prompted us to search for chemokine-related gene enrichment in the transcriptome of PAK2-downregulated HUVECs. GSEA analysis showed strong enrichment of the chemokine production (GO:0032602) and chemokine-mediated signaling pathway (GO:0070098) gene sets in siPAK2 transfected cells (Fig. 4C and S4G). In total, fourteen DEGs coding for chemokines were identified (Fig. 4D). Except for *CCL14*, which was downregulated, all the other chemokines were upregulated in PAK2-depleted HUVECs (Fig. 4D). The differential expression of these chemokines was validated by RT-qPCR (Fig. 4E). Similar to the bulk RNA-seq data, levels of *CXCL10* mRNA were the most upregulated (Fold increase = 37.0) (Fig. 4D, 4E).

Next, to determine if PAK2 kinase activation is necessary for the induction of chemokine expression, we inhibited PAK2 in HUVECs with a pharmacological inhibitor of group I PAKs, FRAX1036 ^29^, and showed that it also caused an augmentation of *CXCL10* mRNA levels (Fig. 4F). Upregulation of other chemokines was also observed in HUVECs following FRAX1036 treatment (Fig. S4H). Inhibition of PAK2 activity by FRAX1036 was confirmed by the reduction in the levels of Ser-141 phosphorylation on PAK2 (Fig. 4F). Importantly, depletion of PAK2 increased the expression of the CXCL10 protein in HUVECs (Fig. 4G). CXCL10 levels were also markedly increased in the cell culture medium of siPAK2-tranfected HUVECs (Fig. 4G). These results suggest that inhibition of PAK2 in ECs increases the immunological response by promoting a pro-inflammatory endothelial state and by stimulating the expression and production of chemokines by ECs, in particular of CXCL10.

### CXCL10 is required for the inhibition of EC sprouting and chemokine expression caused by PAK2 silencing

Next, we examined if CXCL10 expression is necessary for the effects induced by PAK2 silencing in ECs by double-transfecting HUVECs with siRNAs targeting PAK2 and CXCL10 (siPAK2-siCXCL10) (Fig. S5A). First, to assess the contribution of CXCL10 on the anti-angiogenic effects, we used a 3-dimensional EC spheroid-sprouting assay in collagen. VEGF-stimulated sprouting from spheroids of HUVECs was inhibited in siPAK2 transfected cells. While silencing of CXCL10 expression by siRNA (siCXCL10) did not affect EC sprouting from spheroids, it prevented the inhibitory effect of PAK2 downregulation on VEGF-stimulated EC sprouting in double-transfected cells (siPAK2-siCXCL10) (Fig. 5A, B).

**Figure 5.**
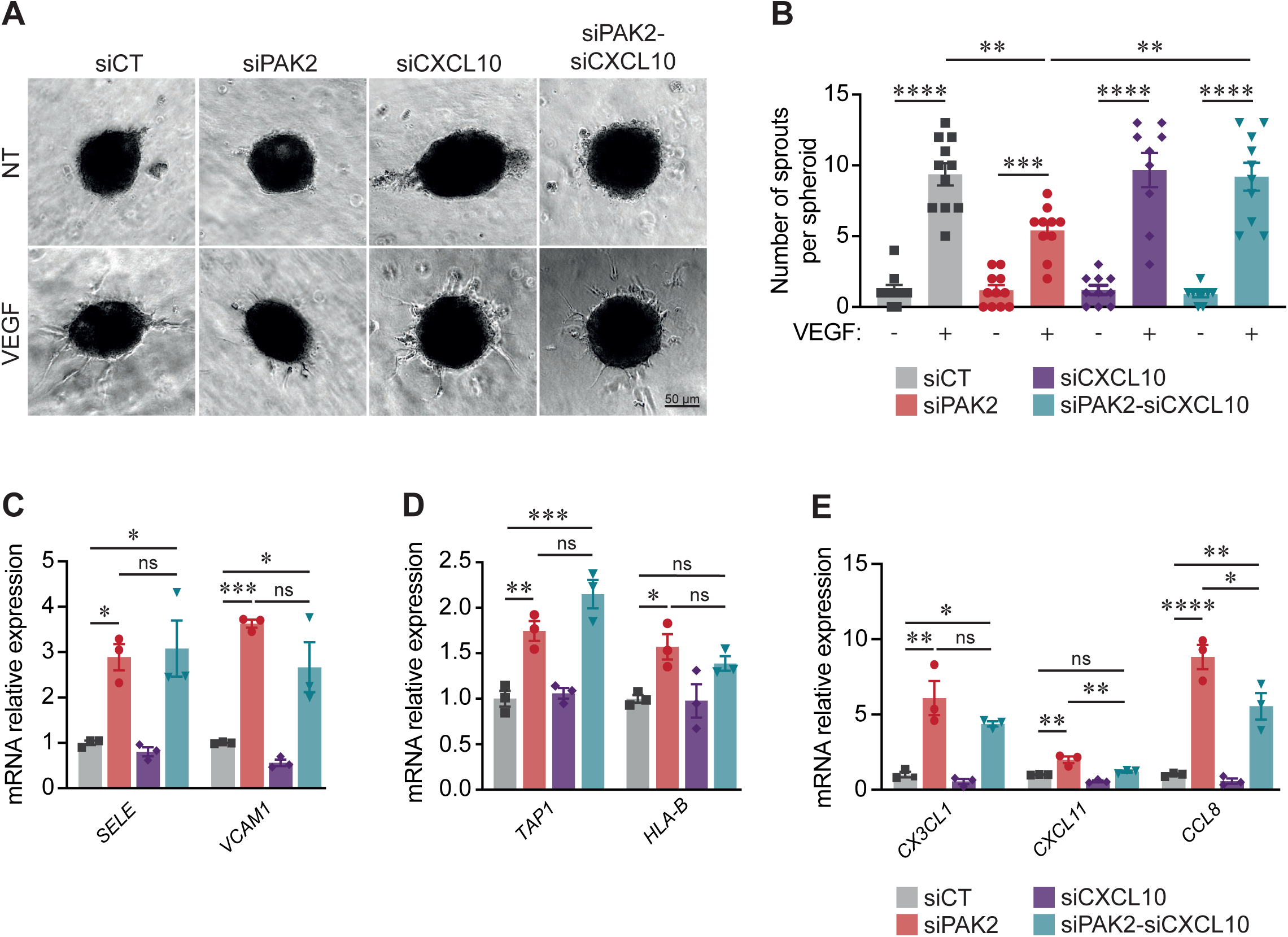
CXCL10 expression is required for anti-angiogenic effects and chemokine expression. **(A)** Representative images from 3-dimensional EC spheroid-sprouting assay generated from HUVECs transfected with control siRNA (siCT), siRNA targeting PAK2 (siPAK2), CXCL10 (siCXCL10) or both (siPAK2-siCXCL10). Spheroid were exposed to VEGF (40 ng/mL) for 24 h after embedding in a collagen matrix. Scale = 50 µm. **(B)** Quantification of the number of capillary-like sprouts per spheroid was measured following VEGF stimulation (n=9-11 spheroids). Data show one of three experiments with similar results. **(C-E)**. Expression of (C) adhesion, (D) antigen processing and presentation, and (E) chemokine genes measured by RT-qPCR, normalized to GAPDH housekeeping gene and expressed as fold change relative to siCT transfected HUVECs for every condition as indicated (n=3). All data are represented as mean ± SEM. *p<0.05, **p<0.01, ***p<0.001, ****p<0.0001.

Then, to assess the contribution of CXCL10 on the immunogenic effects, we examined the involvement of CXCL10 on the expression of genes in PAK2-downregulated ECs that were involved in the enrichment of the above-mentioned GO terms leukocyte adhesion to vascular endothelial cells (GO:0061756), antigen processing and presentation of peptide antigen (GO:0048002) and found among the top DEGs coding for chemokines (Fig. 4D, S4E-F). Increased expression in siPAK2-transfected HUVECs of E-Selectin (*SELE*), Vascular Cell Adhesion Molecule 1 (*VCAM1*), Antigen Peptide Transporter 1 (*TAP1*), Human Leukocyte Antigen-B (*HLA-B*), *CX3CL1*, *CXCL11* and *CCL8* was confirmed by RT-qPCR (Fig. 5C-E). Downregulation of both PAK2 and CXCL10 did not significantly affect the expression of *SELE*, *VCAM1*, *TAP1*, *HLA-B* and *CX3CL1* compared to cells transfected with siPAK2 alone (Fig. 5C-E). However, the expression of *CXCL11* and *CCL8* was significantly reduced in siCXCL10 and siPAK2 transfected ECs compared to PAK2-downregulated cells (Fig. 5E). This suggests that CXCL10 secreted by ECs contributes in an autocrine manner to the inhibition of angiogenesis induced by PAK2 silencing. Similarly, the expression of the chemokines CXCL11 and CCL8 is also controlled by CXCL10 in PAK2-downregulated ECs. In contrast, CXCL10 does not appear to be required for the induction of other inflammatory genes in ECs that are involved in the immunomodulation caused by PAK2 silencing.

### Expression of CXCL10 is increased in KO-PAK2^EC^ tumors

Next, we determined which chemokines were differentially regulated in tumors from CT- and KO-PAK2^EC^ mice. Bulk RNA-seq data of KO-PAK2^EC^ tumors revealed that *Cxcl1*, *Cxcl2* and *Cxcl10* are among DEGs in tumors from KO-PAK2^EC^ mice (Table S1). Similar to the RNA-seq data from PAK2-depleted ECs (Fig. 4), *Cxcl10* had the highest fold change expression (Fold increase = 2.3) in tumors in which endothelial PAK2 expression was suppressed (Fig. 6A). The upregulation of CXCL10 mRNA was confirmed by RT-qPCR in KO-PAK2^EC^ tumors (Fig. S2B). To assess CXCL10 protein levels and its localization relative to tumor blood vessels, CT-PAK2^EC^ and KO-PAK2^EC^ tumor sections were stained for CXCL10 and CD31 by immunofluorescence (Fig. 6B). KO-PAK2^EC^ tumors showed more overall CXCL10-positive signals than CT-PAK2^EC^ tumors, confirming CXCL10 upregulation at the protein level (Fig. 6C). Moreover, the CXCL10-positive signals in KO-PAK2^EC^ tumors were located more in the tumor core than in the periphery (Fig. 6C). However, for both tumor groups CXCL10-positive signals were located in proximity to CD31-positive structures, and no difference in the distances between the CXCL10 signal clusters and tumor ECs was observed in KO-PAK2^EC^ and CT-PAK2^EC^ tumors (Fig. 6D). Overall, these results indicate that CXCL10 production is increased in PAK2-depleted tumors Ecs.

**Figure 6.**
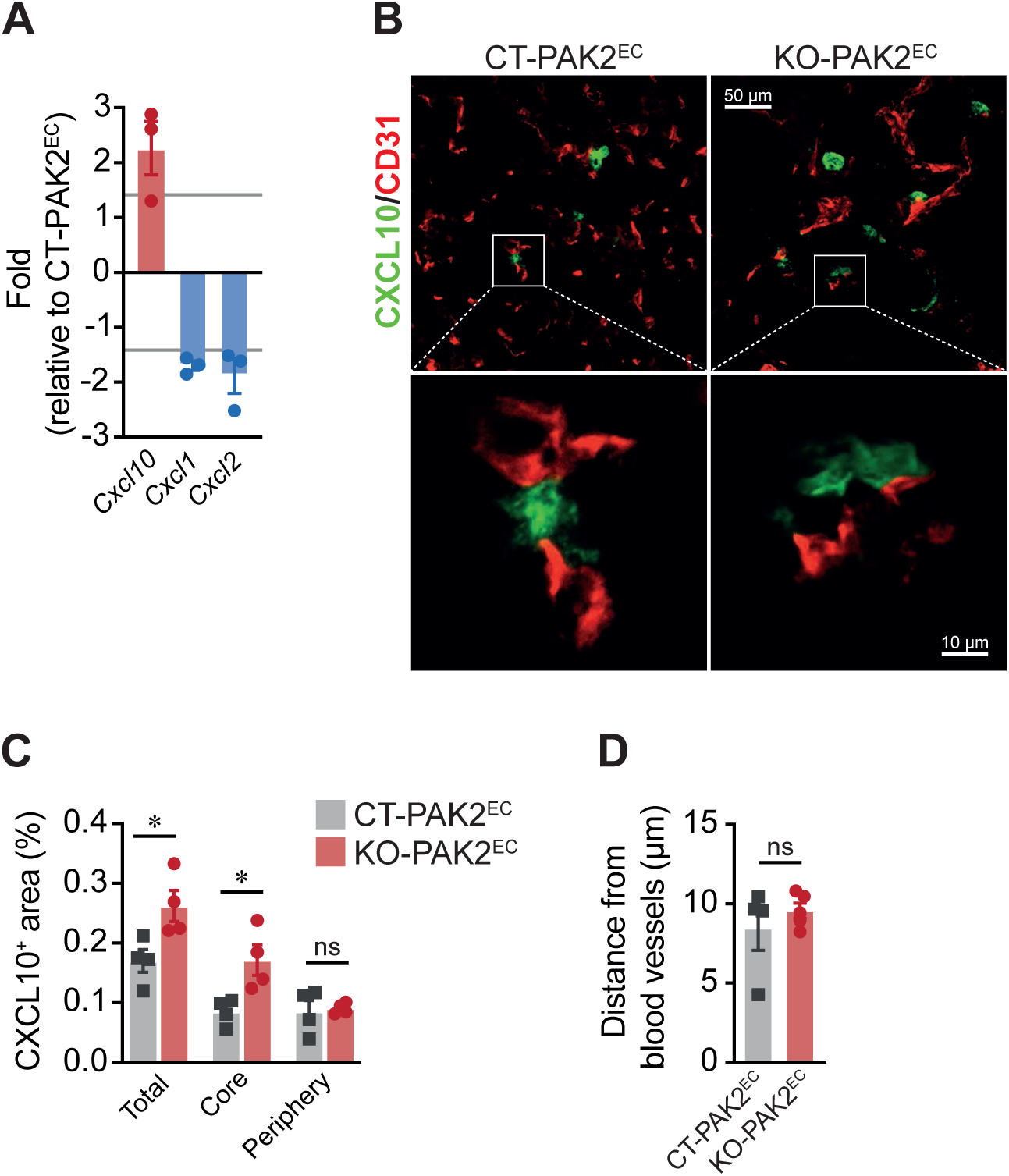
Deletion of PAK2 in tumor ECs enhances CXCL10 expression in tumors. **(A)** Fold change expression of the chemokine DEGs in KO-PAK2^EC^ (n = 3 tumors) relative to CT-PAK2^EC^ tumors (n = 3 tumors) from bulk RNA-seq. **(B)** Visualization of CXCL10^+^ clusters and CD31^+^ vascular structures in tumors. Scale = 50 and 10 μm. **(C)** Quantification of CXCL10^+^ area in total tumor, tumor core and tumor periphery (CXCL10^+^ area/total tumor area) (n=4 tumors). **(D)** Quantification of the distance of CXCL10^+^ clusters from CD31^+^ vascular structures in tumors (n=4-5 tumors). All data are represented as mean ± SEM. *p<0.05.

### Neutralization of CXCL10 abolishes endothelial PAK2 deletion-mediated effects in tumors

To confirm the importance of CXCL10 in the anti-tumoral effects induced by PAK2 deletion in tumor Ecs, neutralizing antibodies against CXCL10 (anti-CXCL10; 50 μg/mouse i.p.) or isotype control IgG2a were administered to CT-PAK2^EC^ or KO-PAK2^EC^ tumor-bearing mice starting on the fourth day of tumor growth and every three days thereafter during the 18-days growth of LLC tumors (Fig. 1A). Similar to Fig. 1B, tumors on control IgG2a-treated KO-PAK2^EC^ mice were significantly smaller than tumors on CT-PAK2^EC^ mice (Fig. 7A). Notably, tumors on KO-PAK2^EC^ mice treated with anti-CXCL10 grew significantly bigger than tumors on control-treated KO-PAK2^EC^ mice, and there was no difference between these and tumors on CT-PAK2^EC^ mice (Fig. 7A). Next, we examined if the reduced angiogenesis and vascular normalization observed in KO-PAK2^EC^ tumors were also affected by CXCL10 blockade. The reduction in CD31-content observed in tumors on KO-PAK2^EC^ mice compared to CT-PAK2^EC^ was abolished by the treatment of KO-PAK2^EC^ mice with anti-CXCL10 (Fig. 7B, C). Furthermore, there was no difference in CD31 staining levels in tumors from anti-CXCL10-treated KO-PAK2^EC^ mice compared to tumors from CT-PAK2^EC^ mice treated with isotype control (Fig. 7B, C). Moreover, CD31-positive vascular structures in these tumors were of smaller diameters (0-10 μm), had decrease pericyte coverage and showed higher fibrinogen leakiness than vascular structures in KO-PAK2^EC^ treated with isotype control as shown by immunostaining indicating that neutralization of CXCL10 in KO-PAK2^EC^ tumors inhibited the vascular normalization induced by *PAK2* deletion in tumor ECs (Fig. 7D-F).

**Figure 7.**
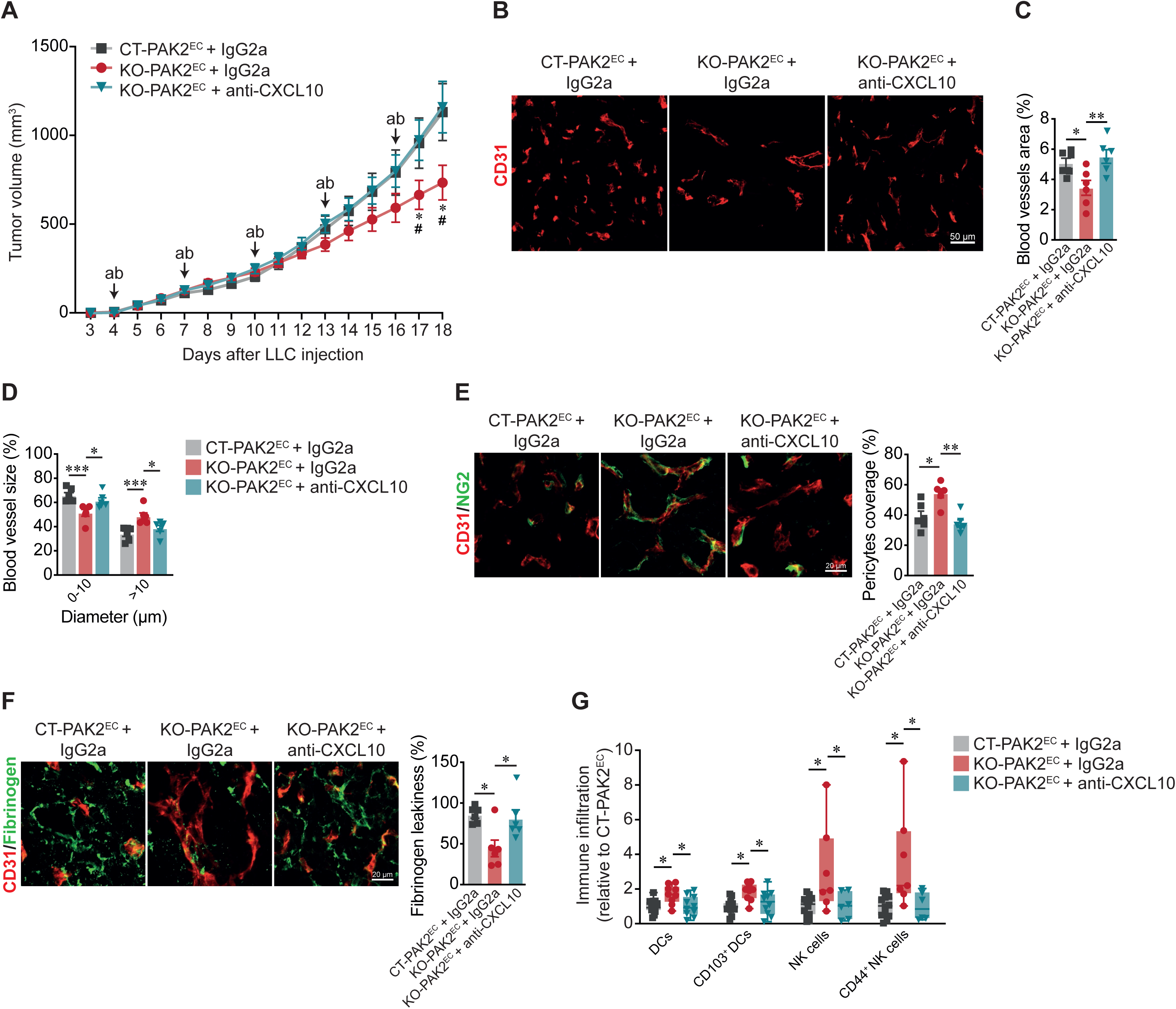
Neutralization of CXCL10 reverses the effects of endothelial-specific deletion of PAK2 on tumor growth, tumor angiogenesis and immune-sensibilization. **(A)** Tumor growth curves of LLC tumors implanted on CT-PAK2^EC^ (n=18 mice) and KO-PAK2^EC^ mice (n=13 mice) treated with control IgG2a isotype, and KO-PAK2^EC^ mice (n=14 mice) treated with CXCL10-neutralizing antibodies (anti-CXCL10, 50 μg i.p.). Arrows indicate time of isotype or antibodies injections. *p<0.05 between CT-PAK2^EC^ control-treated and KO-PAK2^EC^ control-treated mice, #p<0.05 between KO-PAK2^EC^ control-treated and KO-PAK2^EC^ anti-CXCL10 treated mice. **(B)** Representative images of CD31^+^ staining in tumors sections. Scale = 50 μm. **(C)** Quantification of CD31^+^ vascular area in tumors (CD31^+^ area/total area) (n=6 tumors). **(D)** Quantification of diameters of CD31^+^ vascular structures in tumors (n=6 tumors). **(E)** Quantification of NG2^+^ pericyte coverage of blood vessels in tumors (NG2^+^ area/CD31^+^ area) (n=6 tumors). Scale = 20 μm. **(F)** Quantification of Fibrinogen leakage from blood vessels in tumors (Fibrinogen^+^ area/CD31^+^ area) (n=5-6 tumors). Scale = 20 μm. **(G)** Flow cytometry analysis of DCs (CD45^+^I-A/I-E^+^CD11c^+^), migratory CD103^+^ DCs, NK cells (CD45^+^CD8^-^CD4^-^NK1.1^+^) and activated CD44+ NK cells infiltration into KO-PAK2^EC^ control-treated and KO-PAK2^EC^ anti-CXCL10 treated tumors, relative to CT-PAK2^EC^ control-treated tumors (n=6-13 tumors). The number of immune cells for each population was normalized to the number of total live cells for each tumor. Combined data of three independent experiments are shown. (A-F) data are represented as mean ± SEM, and (G) data are represented as median ± minimal to maximal values. *p<0.05, **p<0.01, ***p<0.001.

Finally, the immune cell profiles determined by flow cytometry analyses showed that anti-CXCL10 treatment of KO-PAK2^EC^ mice reduced the amounts of DCs, migratory CD103^+^ DCs, NK cells and of activated CD44^+^ NK cells in tumors (Fig. 7G; Fig. S6A). Importantly, CXCL10 blockade or endothelial PAK2 deletion in mice did not significantly influence the levels of DCs and NK cells in the spleen (Fig. S6B-D). These results confirmed the role of CXCL10 in promoting DCs and NK cells presence in tumors implanted on KO-PAK2^EC^ mice.

Overall, this study demonstrates that CXCL10 produced by Ecs of tumors in mice is central for the reduced angiogenesis, vascular normalization, increased infiltration of DCs and NK cells, and the reduced growth of tumors following the inhibition of PAK2 in Ecs of tumors.

## DISCUSSION

While it was previously known that PAK2 is involved in cellular processes central for angiogenesis ^23–25^, the role of PAK2 in tumor blood vessel formation remained ill-defined. Here, we identify a non-cell autonomous function of PAK2 in Ecs of the TME, which controls tumor angiogenesis and tumor growth. More specifically, PAK2 depletion in tumor Ecs decreased tumor vascularization, normalized remaining tumor blood vessels and increased NK cells and DCs infiltration into tumors, leading to reduced tumor growth. Our results indicate a new role for PAK2 in regulating chemokine expression, especially CXCL10. Neutralization of CXCL10 with blocking antibodies reversed the anti-tumor effects induced by *PAK2* deletion. Thus, we propose PAK2 inhibition in tumor endothelial cells as an actionable anti-angiogenic therapeutic strategy to improve immunotherapy responses.

Although PAK2-deficient animal models have been linked to hemorrhages and increased vascular permeability ^23,30^, we showed that blood vessels in KO-PAK2^EC^ tumors were more stable and less permeable than those found in tumors on control mice. In other settings, PAK activation has also been linked to increased endothelial permeability. For instance, VEGF or thrombotic and inflammatory factors induce PAK activation causing EC contraction and pore formation in the endothelial barrier, ultimately leading to vascular permeability ^31^. PAK activation has also been shown to induce VE-cadherin phosphorylation and internalization, causing a disruption of cell junctions and an increased in permeability to macromolecules in response to VEGF ^32^. Blocking PAK activity *in vivo* also decreased permeability in animal models of acute lung injury and atherosclerosis ^33,34^. Therefore, it has been proposed that PAK proteins play various roles in the maintenance of vascular integrity and in the regulation of endothelial permeability in normal and in pathological conditions ^35^. Our study supports this position as we demonstrate that deletion of PAK2 in ECs results in reduced vascular permeability and in normalization of the tumoral vasculature.

Vascular normalization is generally associated with a boosted immune response in tumors ^36^. Herein, we showed that loss of PAK2 in ECs, in addition to normalizing the tumor vasculature, shifted the TME transcriptome towards a strong innate immune signature and enhanced the infiltration of NK cells, activated CD44^+^ NK cells, DCs and migratory CD103^+^ DCs into tumors. NK cells exert antitumor effects by their cytotoxic activity towards cancer cells and by the secretion of factors orchestrating the immune response in tumors ^37^. There is an active crosstalk between NK cells and DCs, which has been shown to promote their activation and maturation, respectively ^38^. Once mature, DCs adopt migratory properties and traffic to tumor draining lymph nodes to prime T cells against cancer cells ^39^. Intra-tumoral CD103^+^ DCs can also re-stimulate cytotoxic T cells directly within the TME ^40^. Although we did not observe increased T cell infiltration in tumors following PAK2 deletion in tumor ECs, it is possible that T cells found in these tumors are more specific against cancer cells than those found in control tumors due to the increase of DCs in KO-PAK2^EC^ tumors and could also contribute to the reduction of tumor growth along with activated NK cells. Interestingly, presence of NK cells in the TME was shown to correlate with intra-tumoral CD103^+^ DCs and with the responsiveness to anti-PD-1 immunotherapy in mouse and human cancers ^41^. This suggests that PAK2 depletion in tumor ECs could be an interesting strategy to improve the response to immunotherapy. The critical role of the tumor immune microenvironment (TIME) in tumor initiation and response to therapy led to the recognition that a better understanding of this process may reveal new therapeutic targets ^42^. Leveraging our deep RNA-seq data, we uncovered an innate immune component upon PAK2 deletion in tumor ECs, which we identified as DCs and NK cells, thereby providing genetic evidence for the importance of PAK2 in shaping the TIME.

Our results align with a recent study that demonstrated that PAK4 inhibition in ECs lead to vascular normalization and immune sensibilization in glioblastoma ^21^, indicating that PAK2 and PAK4 might have overlapping functions in tumor ECs. We did not find in PAK2-depleted ECs any significant change in PAK4 expression indicating that the effects we observed in KO-PAK2^EC^ tumors are specifically caused by the loss of PAK2. Loss of PAK4 in ECs of glioblastoma resulted in ZEB-1 dependent reprogramming of ECs that affected the expression of cell junction proteins and adhesion molecules that contributed to tumor blood vessel normalization and enhanced T-cell recruitment in tumors. We did not observe an effect of PAK2 loss in ECs on *ZEB1* expression indicating that the ZEB1-dependent reprogramming is specific to PAK4 inhibition. Indeed, we identified that the anti-angiogenic and immunogenic effects induced by the deletion of PAK2 are principally mediated by the expression of the chemokine CXCL10 by ECs. Importantly, we demonstrated that CXCL10 neutralization during tumor growth reversed antitumoral effects mediated by the deletion of PAK2 in tumor ECs in mice. CXCL10 exerts well known immunomodulating and angiostatic effects ^14^. ECs, DCs and NK cells express CXCR3, the specific receptor for CXCL10 ^43^. While CXCL10 induces DC and NK cell chemotaxis ^14^, binding of CXCL10 to CXCR3 on ECs inhibits endothelial migration and promotes apoptosis ^44–46^. In addition, CXCL10 can also inhibit endothelial proliferation independently of CXCR3 via the binding to the glycosaminoglycan expressed on EC surface ^47,48^. Our in vitro data support a cell autonomous angiostatic effect of CXCL10 on tumor ECs since CXCL10 silencing prevented the inhibitory effect of PAK2 downregulation on EC sprouting from spheroids. In contrast, the induction of inflammatory genes in ECs did not appear to be dependent on CXCL10 expression. CXCL10 secreted by PAK2-deleted tumoral ECs appears to mostly induces chemotaxis of immune cells in the tumor microenvironment. Overall, these results demonstrated the importance of CXCL10 in the effects induced by the deletion of PAK2 in ECs of tumors.

There is considerable interest for the development of PAK inhibitors in cancer ^49^. Our findings demonstrate that PAK2 deletion in tumor ECs results in decreased tumor growth by inhibiting tumor angiogenesis in addition to reversing vascular abnormalities and reducing immune suppression through the expression of CXCL10. We provide insights into the essential role of PAK2 in tumor angiogenesis and report a novel role for PAK2 in regulating CXCL10 expression in tumors. Thus, PAK2 inhibition could be a promising anti-angiogenic target and could potentially improve immunotherapy response, leading to significantly better clinical outcome of many cancer patients.

## METHODS

### Mice

All animal studies were approved by the Animal Care Committee of the Université de Montréal in agreement with the guidelines established by the Canadian Council on Animal Care. Pak2^fl/fl^ mice were generously provided by Dr. Jonathan Chernoff (Fox Chase Cancer Center) and *Pdgfb*-iCreER mice were generously provided by Dr. Marcus Fruttiger (University College London). *Pdgfb*-iCreER;PAK2^fl/fl^ were viable and fertile with no obvious phenotypes. CreER induction and endothelial deletion of PAK2 was achieved by subcutaneous implantation of placebo or tamoxifen slow-release pellets (5 mg/21 days) (Innovative Research of America) into mice neck two days prior tumor implantation.

### Cell culture and treatments

Lewis lung carcinoma (LLC) cells obtained from American Type Culture Collection (ATCC) were cultured in Dulbecco modified Eagle medium (DMEM, Thermo Fisher) supplemented with 10% fetal bovine serum (FBS, HyClone, GE Healthcare Life Sciences), 2 mM L-glutamine, 100 U/ml penicillin and 100 μg/ml streptomycin (Thermo Fisher).

Microvascular mouse lung ECs (MLECs) were isolated from the lungs of *Pdgfb*-iCre;Pak2^fl/fl^ mice as we previously described ^50^. MLECs were grown in DMEM:F12 (1:1) medium (Gibco), supplemented with 20% FBS, 0.1 mg/mL heparin (Millipore-Sigma), 100 U/ml penicillin, 100μg/ml streptomycin and 0.05 mg/mL Endothelial Cell Growth Supplement (ECGS, Corning Life Sciences). Magnetic Dynabeads (Thermo Fisher) were conjugated with anti-mouse CD102 (clone 3C4, BD Biosciences) antibody. Beads were added and incubated for 1h at 4°C. MLECs were selected in a magnetic field for 5 minutes, repeated two times, following trypsinization. Isolated MLECs were treated with increasing concentrations (0.1, 5 and 50 μM) of 4-hydroxytamoxifen (OH-tamoxifen) (Millipore-Sigma) for 24 h.

Human umbilical vein ECs (HUVECs) obtained from VEC Technologies were cultured in M199 media supplemented with 20% FBS, 2 mM L-glutamine, 100 U/ml penicillin, 100 µg/ml streptomycin and 0.05 mg/mL ECGS. HUVECs were serum starved in M199 media with 0.5% FBS for 3 h followed by a 7 h treatment with 1 μM group I PAK inhibitor FRAX1036 (Selleck Chemicals).

### Tumor implantation and monitoring

LLC tumor cells were grown to 80-90% confluence on the day of the implantation. Cells were washed, trypsinized and resuspended in serum, L-glutamine and penicillin/streptomycin-free DMEM. LLC cells (1.5x06 cells in 100 μL) were injected subcutaneously in the flank of 6 to 8 weeks old mice. Tumor volume was determined daily or every other day using a caliper and applying the formula (volume = 0.523 × (width)^2^ × (length)) to approximate the volume of a spheroid. When measured every other day, the mean of day before and the day after was used to estimate missing tumor volume. If mice reached ethical end point of experiments before 18 days of tumor growth, tumor volume was estimated using exponential growth equation (y = y_0_ × exp(k × x)) determined from tumor measures collected prior to euthanasia. Tumor-bearing *Pdgfb*-iCreER;PAK2^fl/fl^ mice treated with tamoxifen did develop general edema, with an average weight gain of 40% of the mouse’s initial weight, without affecting overall survival. *Pdgfb*-iCreER;PAK2^fl/fl^ mice treated did not develop visible edema.

### *In vivo* treatments

4 days after tumor cells injection, mice were treated with neutralizing anti-CXCL10 antibodies (50μg/100 μL PBS i.p.) or IgG2a control (50μg/100 μL PBS i.p.) (R&D Systems) every three days for remaining tumor growth time. Tumors were measured daily or every other day as previously described, and harvested after 18 days of tumor growth.

### Immunofluorescence and immunohistochemistry staining and analysis

For immunofluorescence staining, tumors were snap-freeze in liquid nitrogen immediately after resection and embedded in Tissue-Tek OCT (Optimal Cutting Temperature) before frozen sectioning on a microtome-cryostat. Tumor sections (7-25 μm) were fixed with 10% formalin-buffered solution (Thermo Fisher) for 10 minutes. 1% bovine serum albumin (BSA)-PBSTT (PBS, 0.5% Triton-X100, 0.02% Tween) was used for permeabilizing and blocking for 1h at room temperature. Slides were incubated overnight at 4°C with primary antibodies (see Table S4) in 1% BSA-PBS solution. Fluorescent secondary antibodies were incubated in 1% BSA-PBS solution for 1 h at room temperature before being mounted in Fluoromount G (Electron Microscopy Sciences). Acquisitions were performed on an LSM 800 confocal laser-scanning microscope (Zeiss). Quantifications were performed on ImageJ version 2.14.0 software (National Institute of Health) by applying a threshold on the positive signal level and measuring positive area. For immunofluorescence quantification, 3 to 6 fields (20×) per tumor were scored.

For immunohistochemistry staining, tumors were fixed with 10% formalin-buffered solution (Thermo Fisher) for at least 48 h before being embedded in paraffin. Immunohistochemistry staining were performed by the Histology Core Facility of the Institute of Research in Immunology and Cancerology of the Université de Montréal (IRIC). Primary antibodies are listed in Table S4. Acquisitions were performed on an NanoZoomer 2.0-HT slide scanner (Hamamatsu). Vessel diameters were measured by hand using the NDP.view 2 viewing software (Hamamatsu). For immune cell quantifications and localization, tumors regions were determined manually. Tumor core was defined as highly cellular regions, which consists primarily of tumor cells, and tumor periphery was defined as non-cellular stroma-like regions by analysis of the hematoxylin counterstain. For immunohistochemistry quantification, 6 fields (20×) per tumor were scored.

### Tumor mRNA isolation, RNA-seq and analysis

Total mRNA from 3 CT-PAK2^EC^ and 3 KO-PAK2^EC^ average-sized tumors was extracted using TRIZOL reagent (Invitrogen) according to recommended procedures. Total mRNA was cleaned up using RNeasy columns (Qiagen) and on-column Dnase I treatment was performed using the Rnase-Free Dnase set (Qiagen). Tumor mRNA samples were submitted to the Montreal Clinical Research Institute (IRCM) Molecular Biology Core Facility for RNA-sequencing. Sequencing was performed at the Génome Québec Innovation Centre (McGill University) by using the Illumina HiSeq 2000 platform. The quality of the raw reads was assessed with FASTQC version 0.11.8 and combined with MultiQC version 1.3. The reads were aligned to the GRCm38 genome with STAR version 2.5.1b and the raw alignment counts were calculated with featureCounts version 1.6. DESeq2 version 1.24.0 was then used to normalize gene readcounts. A cut-off of >0.5 log2(fold change) and <-0.5 log2(fold change), and an adjusted p-value <0.05 was applied to define differentially expressed genes (DEGs) between KO-PAK2^EC^ and CT-PAK2^EC^ tumors. Bioinformatics analyses were performed using the Database for Annotation, Visualization and Integrated Discovery (DAVID) version v2023q3 (National Institutes of Health) using DEGs, the enrichment of Gene Ontology (GO) categories and Kyoto Encyclopedia of Genes and Genomes (KEGG) pathway were analyzed, and the Gene Set Enrichment Analysis (GSEA) software version 4.1.0 (Broad Institute and University of California San Diego) using all sequenced genes with a base mean over 100 reads.

### Flow cytometry experiments

As we previously described ^51^, mechanical tumor dissociation was done using gentleMACs Dissociator (Miltenyi Biotec) in RPMI media. After centrifugation, red blood cells were lysed with RBC lysis buffer (Gibco) 5 minutes on ice. After PBS wash, cells were marked with FVS-510 viability reagent (BD Bioscience) 15 minutes at room temperature. After 2% FBS/PBS wash, Fc blocking antibodies CD16/CD32 (eBioscience) were incubated for 15 minutes on ice. Brilliant Stain buffer (BD Horizon) and antibodies targeting extracellular proteins were then added and incubated for 30 minutes at room temperature (see Table S4). After 2% FBS/PBS wash, cells were either fixed with 2% paraformaldehyde in 2% FBS/PBS solution or stained with antibodies targeting intracellular proteins using the FOXP3/Transcription Factor Staining Buffer set (eBioscience) according to the manufacturer’s instructions. For compensation settings, cells and eBeads (eBioscience) were stained with every antibody individually. BD LSR Fortessa and DiVa software version 8 (BD Biosciences) was used for the acquisition and flow cytometry analysis was performed using FlowJo software version 10 (BD Biosciences).

### HUVECs transfections

PAK2 small interfering RNA (siRNA) and non-silencing control siRNA were generated by Horizon Discoveries. siCT 5’ AUGAACGUGAAUUGCUCAAUU 3’, siPAK2 5’ UCACCAAACUAGAGCAAAAUU 3’ , and siCXCL10 5’ AAUCCAAGGUCUUUAGAAAUU 3’ sequences were used. HUVECs were transfected with siRNAs Lipofectamine 2000 (Thermo Fisher) according to recommended procedures for 48 h to 96 h.

### HUVECs mRNA isolation, RNA-seq and analysis

Total mRNA from 3 siCT and 3 siPAK2 independent experiments was extracted using TRIZOL reagent (Invitrogen) according to recommended procedures. Total mRNA was cleaned up using RNeasy columns (Qiagen). HUVEC mRNA samples were submitted to the Institute for Research in Immunology and Cancer (IRIC) Genomics Core Facility. Sequencing was performed at IRIC (Université de Montréal) the by using the Illumina NextSeq500 platform. Quality of the raw reads was assessed with MultiQC version 1.11. Sequences were trimmed for sequencing adapters and low quality 3’ bases using Trimmomatic version 0.35 and aligned to the reference human genome version GRCh38 (gene annotation from Gencode version 37, based on Ensembl 103) using STAR version 2.7.1a. Gene expressions were obtained from STAR both as readcounts and TPM values as well as computed using RSEM in order to obtain normalized gene and transcript level expression, in FPKM and TPM values. DESeq2 version 1.30.1 was then used to normalize gene readcounts. Differential gene expression and bioinformatics analyses were performed as previously described for tumor RNA-seq analysis.

### Immunoblotting

HUVECs were solubilized with lysis buffer containing 1% Nonidet P-40, 0.1% sodium dodecyl sulfate (SDS), 0.1% deoxycholic acid, 50 mM Tris–HCl (pH 7.4), 150 mM NaCl, 0.1 mM ethylenediaminetetraacetic acid (EDTA), 0.1 mM ethylene glycol tetraacetic acid (EGTA), 20 mM sodium fluoride, 1 mM sodium pyrophosphate, and 1 mM sodium orthovanadate and Complete mini EDTA free protease inhibitor cocktail (Roche Diagnostics). Soluble proteins were separated by SDS-PAGE, transferred onto a nitrocellulose membrane (Hybond-ECL, GE Healthcare Life Science) and western blotted. Detection and quantification were performed using HRP-coupled secondary antibodies and and an Image Quant LAS4000 chemiluminescence-based detection system (enhanced chemiluminescence) (GE Healthcare Life Science).

## 3-Dimensional EC spheroid-sprouting assay

24 h after siRNA transfection, transfected HUVECs were cultured in M199 containing 0.3% methylcellulose (Sigma-Aldrich), 20% FBS, 2 mM L-glutamine, 100 U/ml penicillin, and 100 µg/ml streptomycin in a U-shaped 96-well plates for 48 h to allow spheroid formation at 37 C, 5% CO2. Spheroid of approximatively 1500 cells were transferred in complete medium containing 50% collagen (Corning) pH 7.4, 5% HBSS 10X (WISENT Inc.), 0.6% methylcellulose in 24-well plates. Spheroid were stimulated with 40 ng/mL VEGF (R&D System), and cultured for 24 h at 37°C, 5% CO2 before fixation in 5% PFA for 20 min at room temperature. Image acquisitions were performed using an inverted Axio-observer Z1 epifluorescent microscope (Zeiss) with a 10× objective. The number of sprouts per spheroid was manually counted using ImageJ 2.14.0v software. Assay was performed in 3 independent experiments with similar results.

### Quantitative RT-qPCT validation of RNA-sequencing data

cDNAs were generated from 1 μg off RNA isolated using the SuperScript II Reverse Transcriptase kit for RT-qPCR (Thermo Fisher) according to the manufacturer’s instructions. cDNA (0.2 μg) was amplified using PowerUP SYBR Green Master mix (Thermo Fisher). Quantitative real-time PCR (RT-qPCR) was performed with MIC qPCR cycler (Bio Molecular Systems). Gene expression analysis was performed using the comparative cycle threshold (ΔCT) method, normalized with the expression of reference gene GAPDH. Primer sequences are listed in Table S5.

### Statistics

Data are represented as the means ± SEM, except for flow cytometry analysis that are represented as box-and-whisker plot showing median ± minimal to maximal values, from at least 3 independent experiments. Two-tailed independent Student’s t tests were used when comparing two groups. Comparison between multiple groups were made using one-way ANOVA or two-way ANOVA followed by post-hoc Tukey’s multiple comparison test among groups. For cytometry analyses, outliers were identified and excluded using ROUT method (Q1%). All statistical calculation were performed using GraphPad Prism software (version 10.0.3, Dotmatics). P-value <0.05 was considered statistically significant.

### Data availability

All RNA-seq data generated in this study will be deposited in the Gene-Expression Omnibus (GEO).

## AUTHORS’ CONTRIBUTIONS

J.C. designed and performed the experiments, analyzed the data, prepared figures, and wrote the manuscript; P.M., C.G., C.D. and Y.E.B. performed experiments, analyzed data and revised the manuscript; M.-A.G. provided key expertise and revised the manuscript; T.H. analyzed data, provided key expertise and revised the manuscript; J.-F.C. obtained funding, designed experiments and revised the manuscript; J.-P.G. obtained funding, designed, and supervised the experiments, analyzed the data, prepared figures, and wrote the manuscript.

## ACKNOWLEDMENTS

We thank Eric Massicotte and Julie Lord for their assistance with flow cytometry acquisitions. This work was funded by operating grants from the Cancer Research Society of Canada to J.-P.G. (25151 and 937718), from the Leukemia and Lymphoma Society of Canada to T.H., and from the Canadian Institutes of Health Research (CIHR) to J.-F.C. (PJT – 178083), T.H. (PJT – 190275) and J.-P.G (PJT – 180376). J.C. is a recipient of a doctoral training studentship from Fonds de Recherche du Québec – Santé (FRQS). J.-F.C. holds the Canada Research Chair Tier-1 in Cellular Signaling and Cancer Metastasis and the Alain Fontaine Chair in cancer research from the IRCM Foundation. J.-P.G. holds the Merck Canada Chair in Pharmacology from Université de Montréal.

