## Supplementary Figures S1-S6 and Legends for "Inhibition of PAK2 in endothelial cells suppresses tumor angiogenesis and promotes immune sensitization through CXCL10"

**Running Title: PAK2 controls tumor angiogenesis and immune evasion**

**Keywords:** PAK2, CXCL10, tumor angiogenesis, tumor immunity, tumor microenvironment

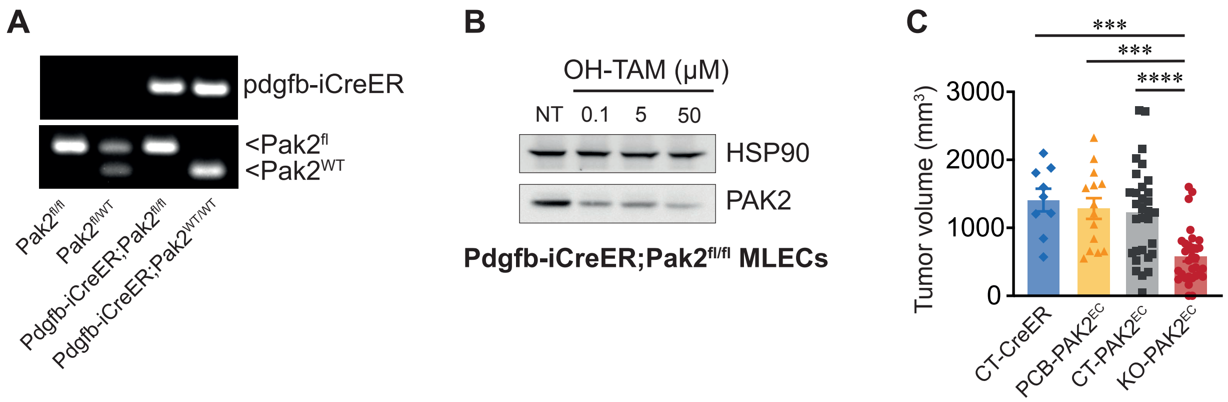

A
pdgfb-iCreER
<Pak2fl
<Pak2WT
Pak2fl/fl
Pak2fl/WT
Pdgfb-iCreER;Pak2fl/fl
Pdgfb-iCreER;Pak2WT/WT
B
OH-TAM (μM)
NT
0.1
5
50
HSP90
PAK2
Pdgfb-iCreER;Pak2fl/fl MLECs
C
***
***
****
Tumor volume (mm3)
3000
2000
1000
0
CT-CreER
PCB-PAK2EC
CT-PAK2EC
KO-PAK2EC

**Figure S1. Pdgfb-iCreER;Pak2$^{fl/fl}$ model induces PAK2 deletion upon tamoxifen treatment responsible for reduced tumor growth. (A)** PCR analysis of mice genotypes. Pdgfb-iCreER;Pak2$^{fl/fl}$ mice were generated by crossing Pak2$^{fl/fl}$ with Pdgfb-iCreER mice. **(B)** Western blot analysis of PAK2 protein expression in mouse lung ECs (MLECs) isolated from Pdgfb-iCreER;Pak2$^{fl/fl}$ mice and treated in vitro with increasing doses of 4-hydroxytamoxifen (OH-tamoxifen) for 24 h. **(C)** Final LLC tumor volume after 18 days of tumor growth. CT-CreER (n=9 mice), PCB-PAK2$^{EC}$ (n=14 mice), CT-PAK2$^{EC}$ (n=31 mice), KO-PAK2$^{EC}$ (n=31 mice). Data are represented as mean ± SEM. ***$p<0.001$, ****$p<0.0001$.

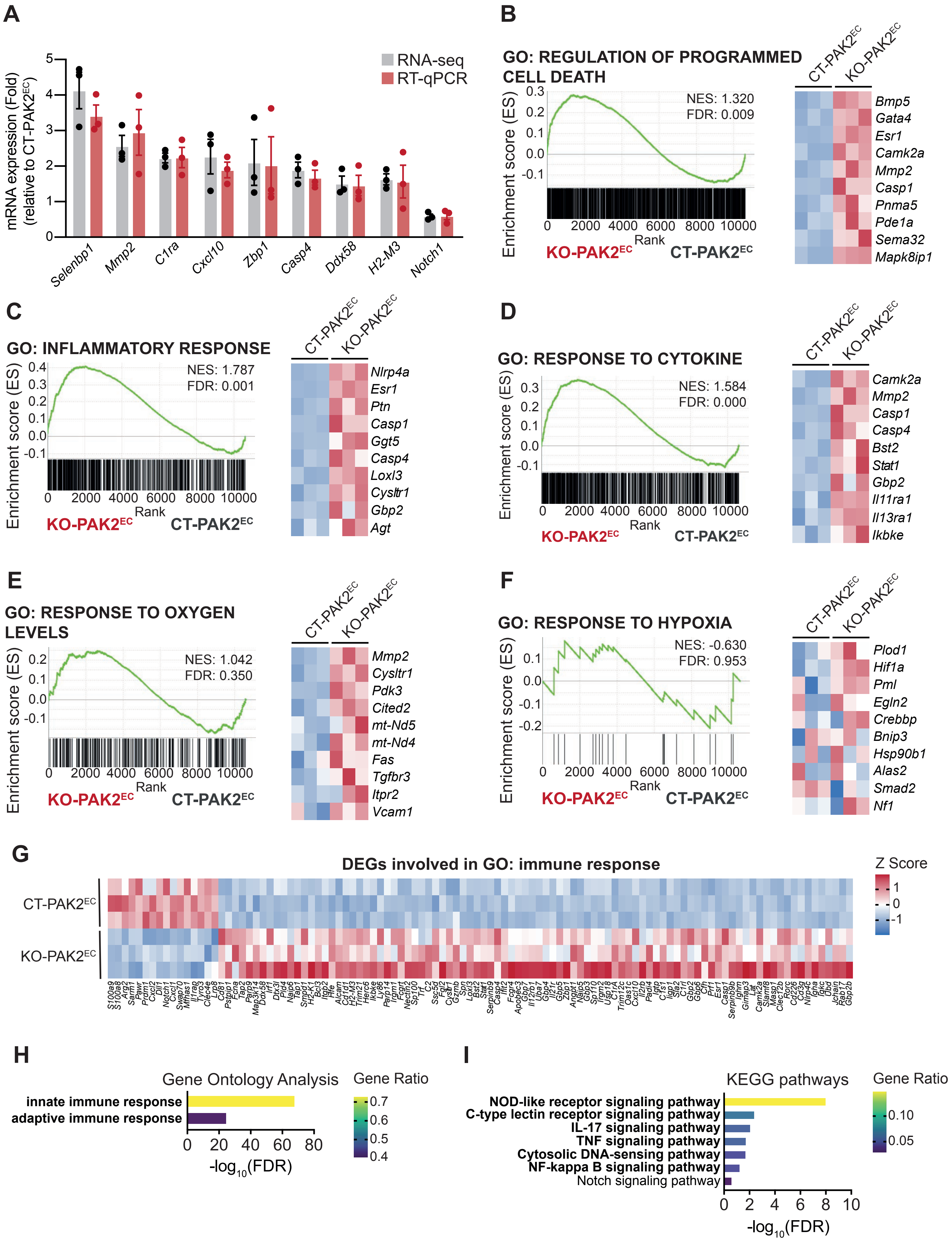
A
mRNA expression (Fold)
(relative to CT-PAK2EC)
RNA-seq
RT-qPCR
Selenbp1
Mmp2
C1ra
Cxcl10
Zbp1
Casp4
Ddx58
H2-M3
Notch1
B
GO: REGULATION OF PROGRAMMED CELL DEATH
NES: 1.320
FDR: 0.009
Enrichment score (ES)
Rank
KO-PAK2EC
CT-PAK2EC
Bmp5
Gata4
Esr1
Camk2a
Mmp2
Casp1
Pnma5
Pde1a
Sema32
Mapk8ip1
C
GO: INFLAMMATORY RESPONSE
NES: 1.787
FDR: 0.001
Nlrp4a
Esr1
Ptn
Casp1
Ggt5
Casp4
Loxl3
Cysltr1
Gbp2
Agt
D
GO: RESPONSE TO CYTOKINE
NES: 1.584
FDR: 0.000
Camk2a
Mmp2
Casp1
Casp4
Bst2
Stat1
Gbp2
Il11ra1
Il13ra1
Ikbke
E
GO: RESPONSE TO OXYGEN LEVELS
NES: 1.042
FDR: 0.350
Mmp2
Cysltr1
Pdk3
Cited2
mt-Nd5
mt-Nd4
Fas
Tgfbr3
Itpr2
Vcam1
F
GO: RESPONSE TO HYPOXIA
NES: -0.630
FDR: 0.953
Plod1
Hif1a
Pml
Egln2
Crebbp
Bnip3
Hsp90b1
Alas2
Smad2
Nf1
G
DEGs involved in GO: immune response
Z Score
CT-PAK2EC
KO-PAK2EC
H
Gene Ontology Analysis
Gene Ratio
innate immune response
adaptive immune response
-log10(FDR)
I
KEGG pathways
Gene Ratio
NOD-like receptor signaling pathway
C-type lectin receptor signaling pathway
IL-17 signaling pathway
TNF signaling pathway
Cytosolic DNA-sensing pathway
NF-kappa B signaling pathway
Notch signaling pathway
-log10(FDR)

**Figure S2. Deletion of PAK2 in ECs of tumors affects the microenvironment transcriptome.** **(A)** RT-qPCR validation of bulk RNA-seq of selected genes. RT-qPCR gene expression analysis was performed using the comparative cycle threshold method, normalized using reference gene GAPDH. Data are represented as mean ± SEM. **(B-F)** GSEA of GO terms (B) regulation of programmed cell death (GO:0043067), (C) inflammatory response (GO:0006954), (D) response to cytokine (GO:0034097), (E) and response to oxygen levels (GO:0070482) (F) response to hypoxia (GO:0001666) in KO-PAK2$^{EC}$ compared to CT-PAK2$^{EC}$ tumors. Heatmaps represent the top 10 genes contributing to the normalized enrichment score (NES). The scale for Z scores in the heatmaps of panels B-F is shown in panel G. **(G)** Heatmap of the DEGs involved in the GO term immune response (GO:0006955). **(H)** Bar chart of the GO terms innate immune response (GO:0045087) and adaptative immune response (GO:0002250) of DEGs involved in the immune response GO term. The x-axis indicates the statistical significance of the enrichment (-$\log_{10}$(FDR)). Color-coding indicates the gene ratio, referencing to the ratio of DEGs annotated in the given GO term to all genes annotated in the GO term. **(I)** Bar graph of significant KEGG pathways of DEGs involved in the immune response GO term. The x-axis indicates the statistical significance of the enrichment (-$\log_{10}$(FDR)). Color-coding indicates the gene ratio, referencing to the ratio of DEGs annotated in the given GO term to all genes annotated in the GO term.

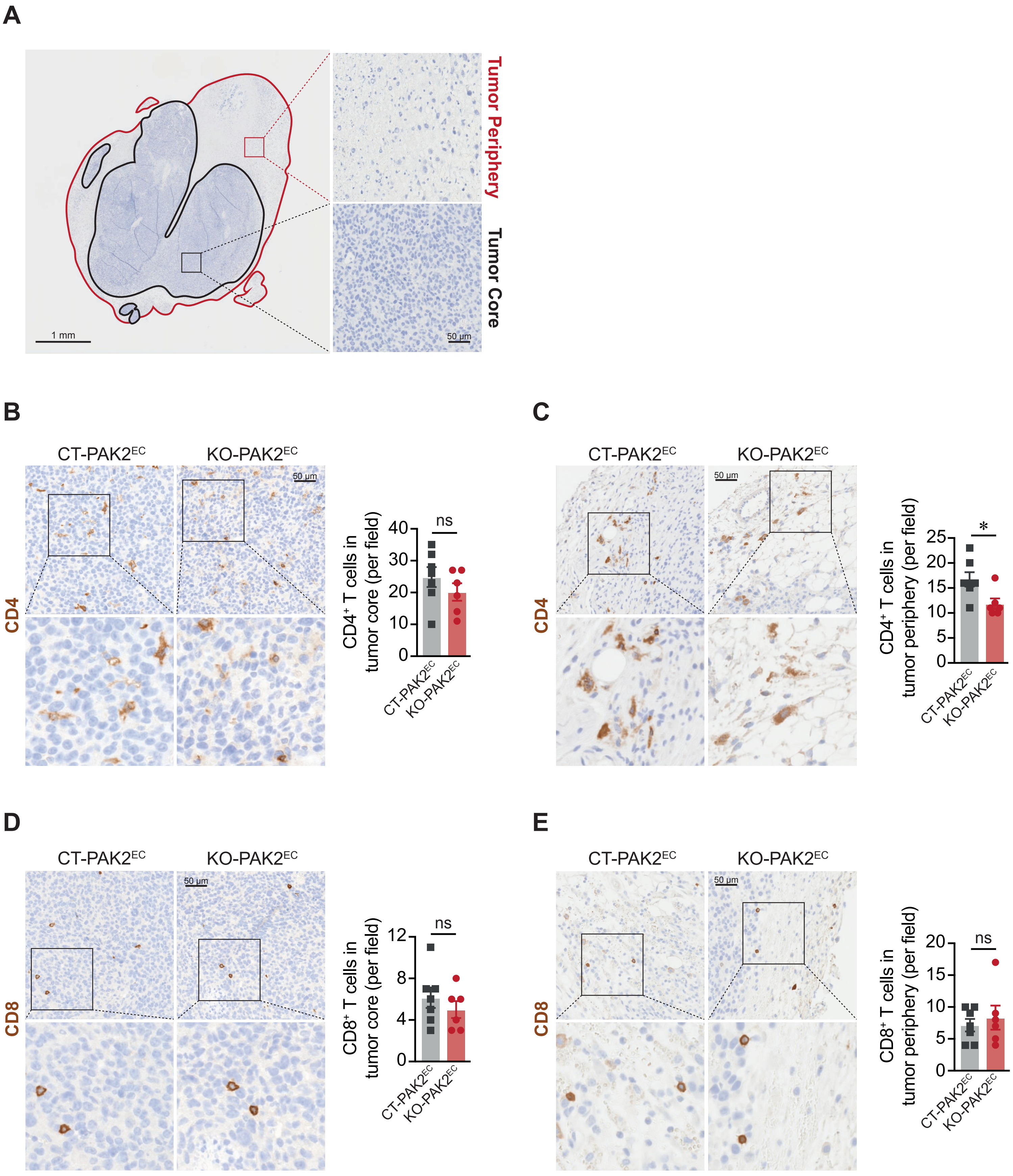
A
Tumor Periphery
Tumor Core
1 mm
50 µm
B
CT-PAK2EC
KO-PAK2EC
50 µm
CD4
CD4+ T cells in tumor core (per field)
ns
C
CT-PAK2EC
KO-PAK2EC
50 µm
CD4
CD4+ T cells in tumor periphery (per field)
*
D
CT-PAK2EC
KO-PAK2EC
50 µm
CD8
CD8+ T cells in tumor core (per field)
ns
E
CT-PAK2EC
KO-PAK2EC
50 µm
CD8
CD8+ T cells in tumor periphery (per field)
ns

**Figure S3. Tumor endothelial-specific deletion of PAK2 does not affect T cells infiltration in the core of tumors. (A)** Visualization of tumor core and tumor periphery in tumors based on hematoxylin counterstain. Tumor core was defined as highly cellular regions, and tumor periphery was noncellular stroma-like regions. Scale = 1 mm and 50 µm. **(B-E)** Immunohistochemistry analysis of KO-PAK2$^{EC}$ and CT-PAK2$^{EC}$ tumors. Quantification of CD4+ T cells in the (B) core regions and (C) periphery of tumors (n=6-7). Quantifications of CD8+ T cells in the (D) core regions and (E) periphery of tumors (n=6-7). All data are represented as mean ± SEM. *$p<0.05$.

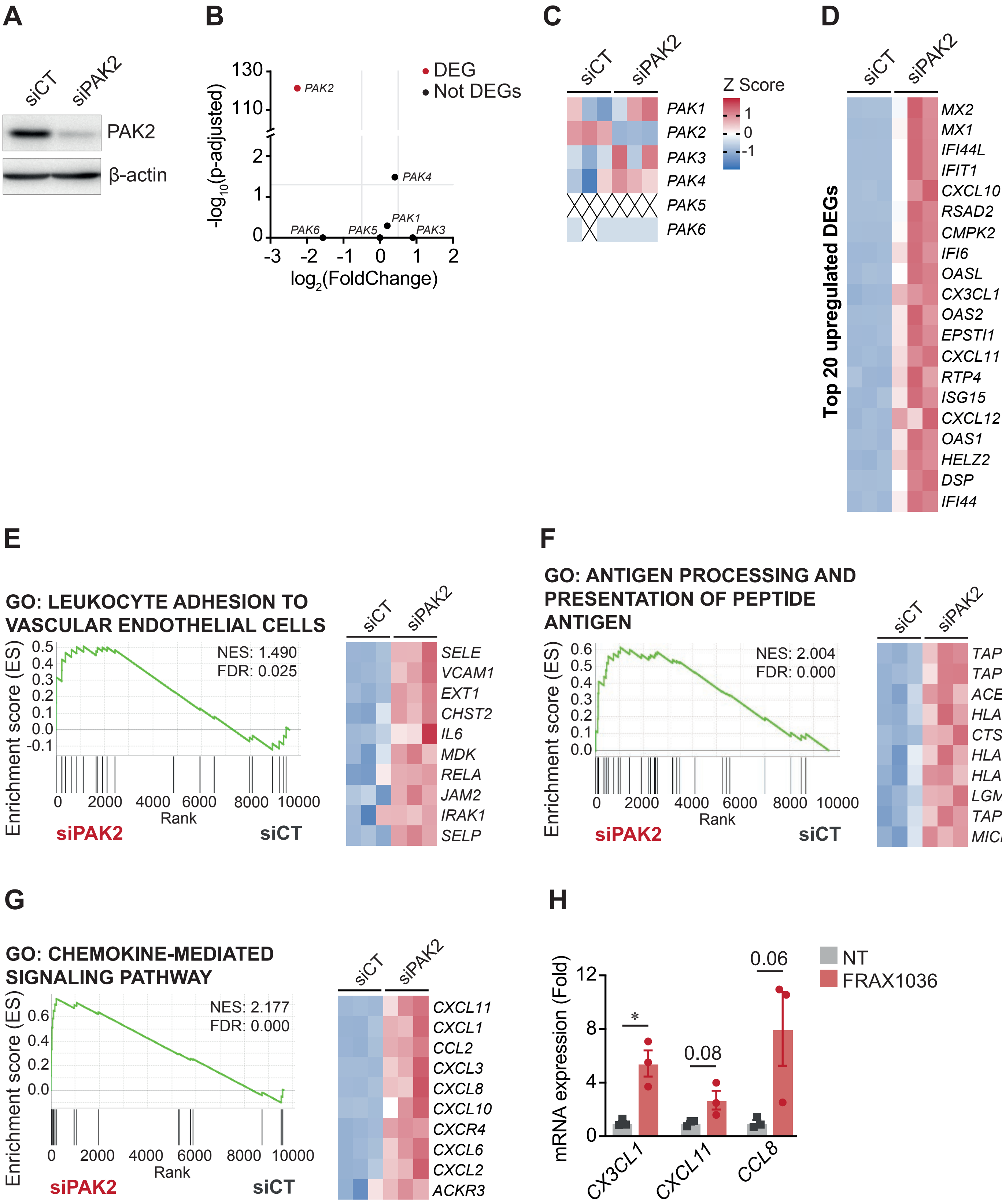

A
siCT
siPAK2
PAK2
β-actin
B
DEG
Not DEGs
-log10(p-adjusted)
log2(FoldChange)
PAK2
PAK4
PAK1
PAK6
PAK5
PAK3
C
Z Score
PAK1
PAK2
PAK3
PAK4
PAK5
PAK6
D
Top 20 upregulated DEGs
MX2
MX1
IFI44L
IFIT1
CXCL10
RSAD2
CMPK2
IFI6
OASL
CX3CL1
OAS2
EPSTI1
CXCL11
RTP4
ISG15
CXCL12
OAS1
HELZ2
DSP
IFI44
E
GO: LEUKOCYTE ADHESION TO VASCULAR ENDOTHELIAL CELLS
NES: 1.490
FDR: 0.025
Enrichment score (ES)
Rank
siPAK2
siCT
SELE
VCAM1
EXT1
CHST2
IL6
MDK
RELA
JAM2
IRAK1
SELP
F
GO: ANTIGEN PROCESSING AND PRESENTATION OF PEPTIDE ANTIGEN
NES: 2.004
FDR: 0.000
Enrichment score (ES)
Rank
siPAK2
siCT
TAP1
TAP2
ACE
HLA-B
CTSS
HLA-C
HLA-A
LGMN
TAPBP
MICB
G
GO: CHEMOKINE-MEDIATED SIGNALING PATHWAY
NES: 2.177
FDR: 0.000
Enrichment score (ES)
Rank
siPAK2
siCT
CXCL11
CXCL1
CCL2
CXCL3
CXCL8
CXCL10
CXCR4
CXCL6
CXCL2
ACKR3
H
mRNA expression (Fold)
NT
FRAX1036
*
0.08
0.06
CX3CL1
CXCL11
CCL8

**Figure S4. Specific downregulation of PAK2 in ECs enhances pro-inflammatory gene expression and chemokine pathways. (A)** Depletion of PAK2 monitored by Western blot. β-actin was used as a loading control. **(B)** Volcano plot showing PAK genes expression in siPAK2 (n=3) compared to siCT (n=3). A threshold of a $log_2$(Fold Change) >0.5 and <-0.5, and a p-adjusted value <0.05 was used to define DEG (red). **(C)** Heatmap of PAK genes expression in siCT and siPAK2 HUVECs. **(D)** Heatmap of the top 20 upregulated DEGs in siPAK2 compared to siCT transfected HUVECs. **(E-G)** GSEA of GO terms (E) leukocyte adhesion to vascular endothelial cells (GO:0061756), (F) antigen processing and presentation of peptide antigen (GO:0048002) and (G) chemokine-mediated signaling pathway (GO:0070098). Heatmaps represent the top 10 genes contributing to the normalized enrichment score (NES). The scale for Z scores in the heatmaps of panels D-G is shown in panel C. **(H)** Expression of selected chemokine genes measured by RT-qPCR, normalized to GAPDH housekeeping gene and expressed as fold change of FRAX1036 treated HUVECs (n=3) relative to non-treated HUVECs (n=3). HUVECs were treated with FRAX1036 (1 μM) or M199 serum-free media for 7 h.

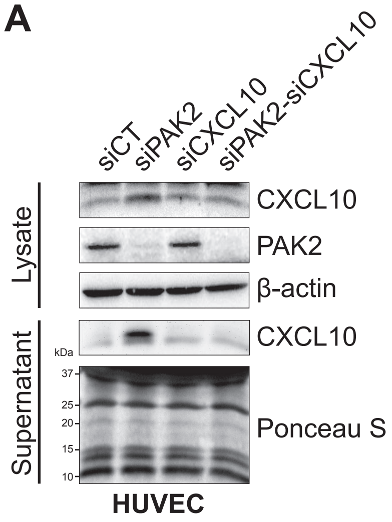
A
siCT
siPAK2
siCXCL10
siPAK2-siCXCL10
Lysate
CXCL10
PAK2
β-actin
Supernatant
CXCL10
kDa
37
25
20
15
10
Ponceau S
HUVEC

**Figure S5. Efficient double transfection of HUVECs with siRNAs targeting PAK2 and CXCL10. (A)** Depletion of PAK2 and CXCL10 monitored by Western blot in cell lysate and/or cell supernatant. β-actin and Ponceau S staining served as loading control for cell lysate and cell supernatant respectively.

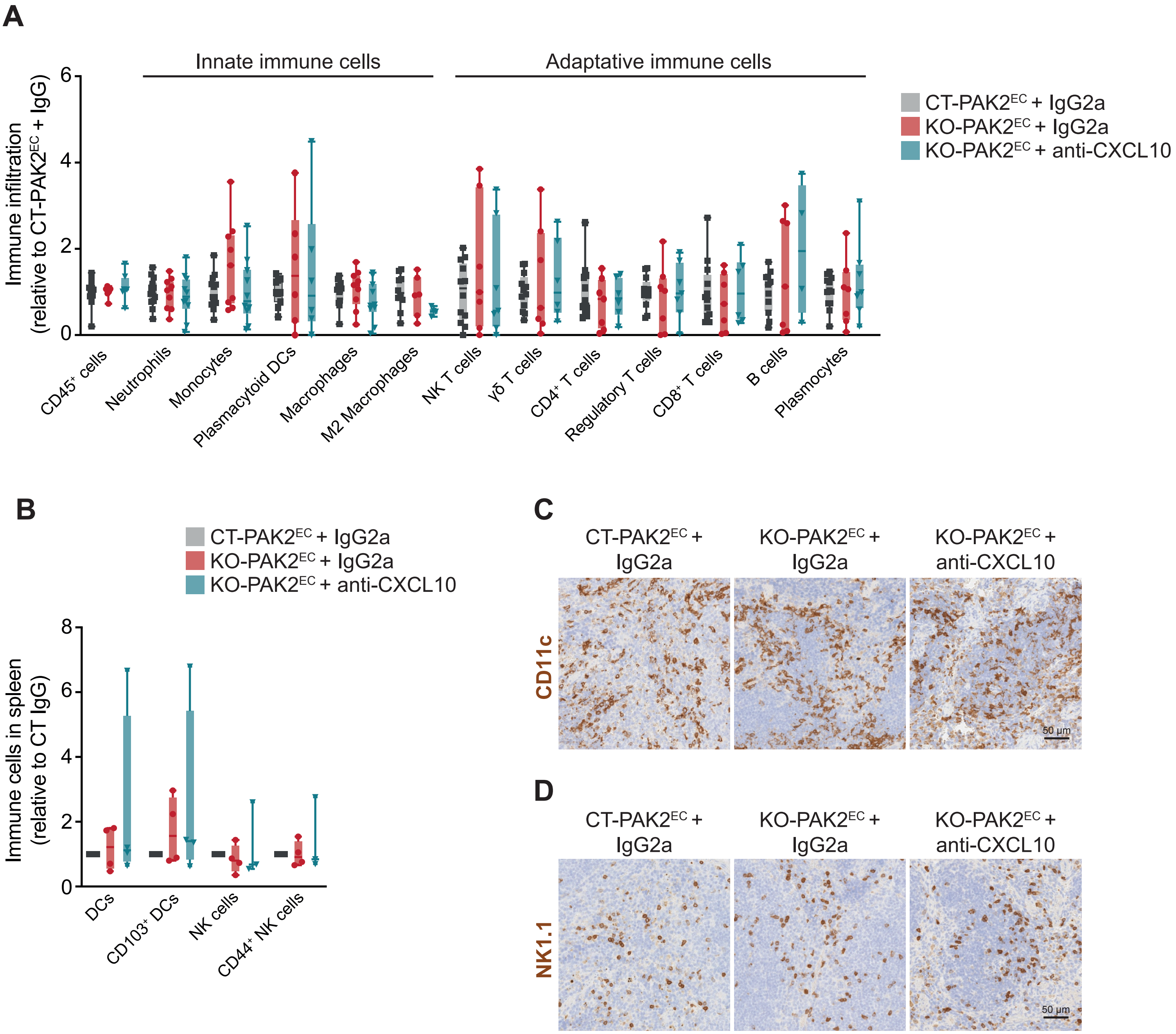
A
Innate immune cells
Adaptative immune cells
Immune infiltration
(relative to CT-PAK2EC + IgG)
6
4
2
0
CT-PAK2EC + IgG2a
KO-PAK2EC + IgG2a
KO-PAK2EC + anti-CXCL10
CD45+ cells
Neutrophils
Monocytes
Plasmacytoid DCs
Macrophages
M2 Macrophages
NK T cells
γδ T cells
CD4+ T cells
Regulatory T cells
CD8+ T cells
B cells
Plasmocytes
B
CT-PAK2EC + IgG2a
KO-PAK2EC + IgG2a
KO-PAK2EC + anti-CXCL10
Immune cells in spleen
(relative to CT IgG)
8
6
4
2
0
DCs
CD103+ DCs
NK cells
CD44+ NK cells
C
CT-PAK2EC + IgG2a
KO-PAK2EC + IgG2a
KO-PAK2EC + anti-CXCL10
CD11c
50 µm
D
CT-PAK2EC + IgG2a
KO-PAK2EC + IgG2a
KO-PAK2EC + anti-CXCL10
NK1.1
50 µm

**Figure S6. Effects of CXCL10 neutralization on immune populations. (A)** Flow cytometry analysis of immune cell infiltration into KO-PAK2$^{EC}$ control-treated and KO-PAK2$^{EC}$ anti-CXCL10 treated tumors, relative to CT-PAK2$^{EC}$ control-treated tumors (n=4-13 tumors). Neutrophils (CD45$^{+}$Ly6G$^{+}$CD11b$^{+}$), monocytes (CD45$^{+}$Ly6C$^{+}$CD11b$^{+}$), plasmacytoid DCs (CD45$^{+}$Ly6C$^{+}$CD11b$^{-}$CD11c$^{+}$CD317$^{+}$), macrophages (CD45$^{+}$F4/80$^{+}$CD11b$^{+}$), M2 macrophages (CD45$^{+}$F4/80$^{+}$CD11b$^{+}$CD206$^{+}$), NK T cells (CD45$^{+}$NK1.1$^{+}$CD3e$^{+}$), γδ T cells (CD45$^{+}$CD3e$^{+}$γδTCR$^{+}$), CD4$^{+}$ T cells (CD45$^{+}$CD3e$^{+}$CD4$^{+}$), regulatory T cells (CD45$^{+}$CD3e$^{+}$CD4$^{+}$CD25$^{+}$FoxP3$^{+}$), CD8$^{+}$ T cells (CD45$^{+}$CD3e$^{+}$CD8$^{+}$), B cells (CD45$^{+}$B220$^{+}$CD19$^{+}$), plasmocytes (CD45$^{+}$B220$^{+}$). The number of immune cells for each population was normalized to the number of total live cells. Combined data of three independent experiments are shown and represented as median ± minimal to maximal values. **(B)** Flow cytometry analysis of DCs (CD45$^{+}$I-A/I-E$^{+}$CD11c$^{+}$), migratory CD103$^{+}$ DCs, NK cells (CD45$^{+}$CD8$^{-}$CD4$^{-}$NK1.1$^{+}$) and CD44$^{+}$ NK cells presence in spleen of KO-PAK2$^{EC}$ control-treated and KO-PAK2$^{EC}$ anti-CXCL10 treated tumors, relative to CT-PAK2$^{EC}$ control-treated tumors (n=3-4 spleens). The number of immune cells for each population was normalized to the number of total live cells. Combined data of three independent experiments are shown and represented as median ± minimal to maximal values. **(C-D)** Representative images of immunochemistry staining of (C) DCs (CD11c$^{+}$) and (D) NK cells (NK1.1$^{+}$) in spleen of CT-PAK2$^{EC}$ control-treated mice, KO-PAK2$^{EC}$ control-treated and KO-PAK2$^{EC}$ anti-CXCL10 treated mice.

**Table S1. Differentially expressed genes (DEGs) between KO-PAK2$^{EC}$ tumors and CT-PAK2$^{EC}$ (.XLSX)**

**Table S2. List of GO annotations based centered on biological processes of the differentially expressed genes (DEGs) in KO-PAK2$^{EC}$ tumors (.XLSX)**

**Table S3. Differentially expressed genes (DEGs) between siPAK2 and siCT transfected HUVECs (.XLSX)**

**Table S4. List of primary antibodies.**

| Antibodies | Source | Application |
|---|---|---|
| CD31 | Biocare Medical 902-303-092017 | Immunofluorescence |
| PAK2 | Invitrogen PA5-20254 | Immunofluorescence |
| NG2 | Millipore AB5320 | Immunofluorescence |
| Fibrinogen | Dako A0080 | Immunofluorescence |
| CXCL10 | Cell Signaling 14969 | Immunofluorescence, Immunoblot |
| CD11c | Cell Signaling 97858 | Immunohistochemistry |
| NK1.1(Klrb1c/CD161c) | Cell Signaling 39197 | Immunohistochemistry |
| CD8 | Cell Signaling 98941S | Immunohistochemistry |
| CD4 | Abcam ab183685 | Immunohistochemistry |
| CD8a PerCP-Cy5.5 | BD Biosciences 561109 | Flow cytometry |
| CD25 BB515 | BD Biosciences 564458 | Flow cytometry |
| CD45 APC-Cy7 | BD Biosciences 561037 | Flow cytometry |
| CD19 APC-R700 | BD Biosciences 565473 | Flow cytometry |
| CD4 APC | BD Biosciences 561091 | Flow cytometry |
| CD3e BV786 | BD Biosciences 564379 | Flow cytometry |
| CD44 BV711 | BD Biosciences 563971 | Flow cytometry |
| I-A/I-E BV650 | BD Biosciences 563415 | Flow cytometry |
| B220 PE-CF594 | BD Biosciences 562313 | Flow cytometry |
| NK1.1 PE-Cy7 | BD Biosciences 562062 | Flow cytometry |
| γδ TCR PE-Cy7 | BD Biosciences 561997 | Flow cytometry |
| FoxP3 PE | eBioscience 48-5773-80 | Flow cytometry |
| CD11b PerCP-Cy5.5 | BD Biosciences 561114 | Flow cytometry |
| Ly6G AF700 | BD Biosciences 561236 | Flow cytometry |
| CD206 AF647 | BD Biosciences 565250 | Flow cytometry |
| CD103 BV786 | BD Biosciences 564322 | Flow cytometry |
| F4/80 BV421 | BD Biosciences 565411 | Flow cytometry |
| CD11c PE-Cy7 | BD Biosciences 561022 | Flow cytometry |

| | | |
|---|---|---|
| Ly6C PE-CF594 | BD Biosciences 562728 | Flow cytometry |
| CD317 PE | eBioscience 12-3172-81 | Flow cytometry |
| CD19 FITC | BioLegend 115505 | Flow cytometry |
| CD3e FITC | BioLegend 100305 | Flow cytometry |
| NK1.1 FITC | BioLegend 108705 | Flow cytometry |
| PAK2 | Cell Signaling 2608 | Immunoblot |
| p-PAK2 (ser141) | Cell Signaling 2606 | Immunoblot |
| HSP90 | BD Biosciences 610419 | Immunoblot |
| β-actine | Cell Signaling 3700S | Immunoblot |
| CXCL10 | R&D System MAB466 | Neutralization |
| IgG2a | R&D System MAB006 | Neutralization |

**Table S5. List of primer sequences used for RT-qPCR quantifications.**

| **Gene** | **Primers** | **Species** |
|---|---|---|
| *Selenbp11* | GAAGTCATGGTCAGCACCTT<br>CTTCTCCCATGTCCCTTTCAC | Mouse |
| *Mmp2* | CCAGATGTGGCCAACTACAA<br>GGTCAGGTGTGTAACCAATGA | Mouse |
| *C1ra* | CTTGAATACCCTCAGCCCTATC<br>TCACCCAAGTTCTTCCCATTAG | Mouse |
| *Cxcl10* | GGCCATAGGGAAGCTTGAAA<br>CAGACATCTCTGCTCATCATTCT | Mouse |
| *Zbp1* | CTCCTGCAATCCCTGAGAAC<br>GTCATAGCTCAGAAGGTGCTTAT | Mouse |
| *Casp4* | GAAAGAGGAGCTTACAGCAGAG<br>TGAGACATTAGCACCAGGAATG | Mouse |
| *Ddx58* | CAGAACTGGAACAGGTCGTTTA<br>GCTTCTCTGTCTCCTTCATCAG | Mouse |
| *H2-M3* | GCCAGCAGGAACAGTGATTA<br>AGGAGACACAGGTATGGAGAG | Mouse |
| *Notch1* | GATGGACGACAATCAGAACGA<br>ATCACTCAGGTCAGGGAGAA | Mouse |
| *Gapdh* | AACTTTGGCATTGTGGAAGG<br>ACACATTGGGGGTAGGAACA | Mouse |
| *CXCL10* | CCATGAATCAAACTGCCATTCT<br>ACTAATGCTGATGCAGGTACA | Human |
| *CX3CL1* | CTTACCAGCAGAGCACCTTAG<br>GCTCTGCCCATTTCCATTTG | Human |
| *CXCL11* | CAACGATGCCTAAATCCCAAATC<br>GTCCTTTCACCCACCTTTCA | Human |
| *CXCL12* | AGAGCCTGAGGGATCTTTACT<br>GGCAGGTACATCCAAGTTCTAC | Human |
| *CCL8* | TGGAGAGCTACACAAGAATCAC<br>AGGCTCATGGCTTCAGATTT | Human |
| *CXCL8* | GGACAAGAGCCAGGAAGAAA<br>GGGTGGAAAGGTTTGGAGTAT | Human |
| *CXCL1* | ACTCAAGAATGGGCGGAAAG<br>CTTCTCCTAAGCGATGCTCAAA | Human |
| *CCL2* | CTGTGCCTGCTGCTCATA<br>CTGTGCCTGCTGCTCATA | Human |
| *CXCL3* | CTTCTCGCACAGCTTCCC | Human |

| | | |
|---|---|---|
| | AGACAAGCTTTCTTCCCATTCT | |
| *CXCL6* | GCTGCGTTGCACTTGTTTAC<br>TCCAGACAAACTTGCTTCCC | Human |
| *CXCL5* | CCTGAAGAACGGGAAGGAAA<br>CTGCTGAAGACTGGGAAACT | Human |
| *CXCL2* | GCAGGGAATTCACCTCAAGAA<br>TCCTTCTGGTCAGTTGGATTTG | Human |
| *CCL14* | AACAGCCAGTGCTCCAAG<br>TGAGAGTTAGCGGTGGGT | Human |
| *SELE* | GGGAATTGGGACAACGAGAA<br>GGTGAAGTTGCAGGATGATTTG | Human |
| *VCAM1* | ACACAGGTGGGACACAAATAA<br>AATGAGACGGAGTCACCAATC | Human |
| *TAP1* | GGGTGACGGGATCTATAACAAC<br>TCCTCTGTTACCCGAGACAT | Human |
| *HLA-B* | GTCCTAGCAGTTGTGGTCATC<br>TCTCAGTCCCTCACAAGACA | Human |
| *GAPDH* | CCACTCCTCCACCTTTGAC<br>ACCCTGTTGCTGTAGCCA | Human |
